## Supplemental Figures and Tables for "*Casilio-ME*: Enhanced CRISPR-based DNA demethylation by RNA-guided coupling methylcytosine oxidation and DNA repair pathways"

### Supplementary Figure 1

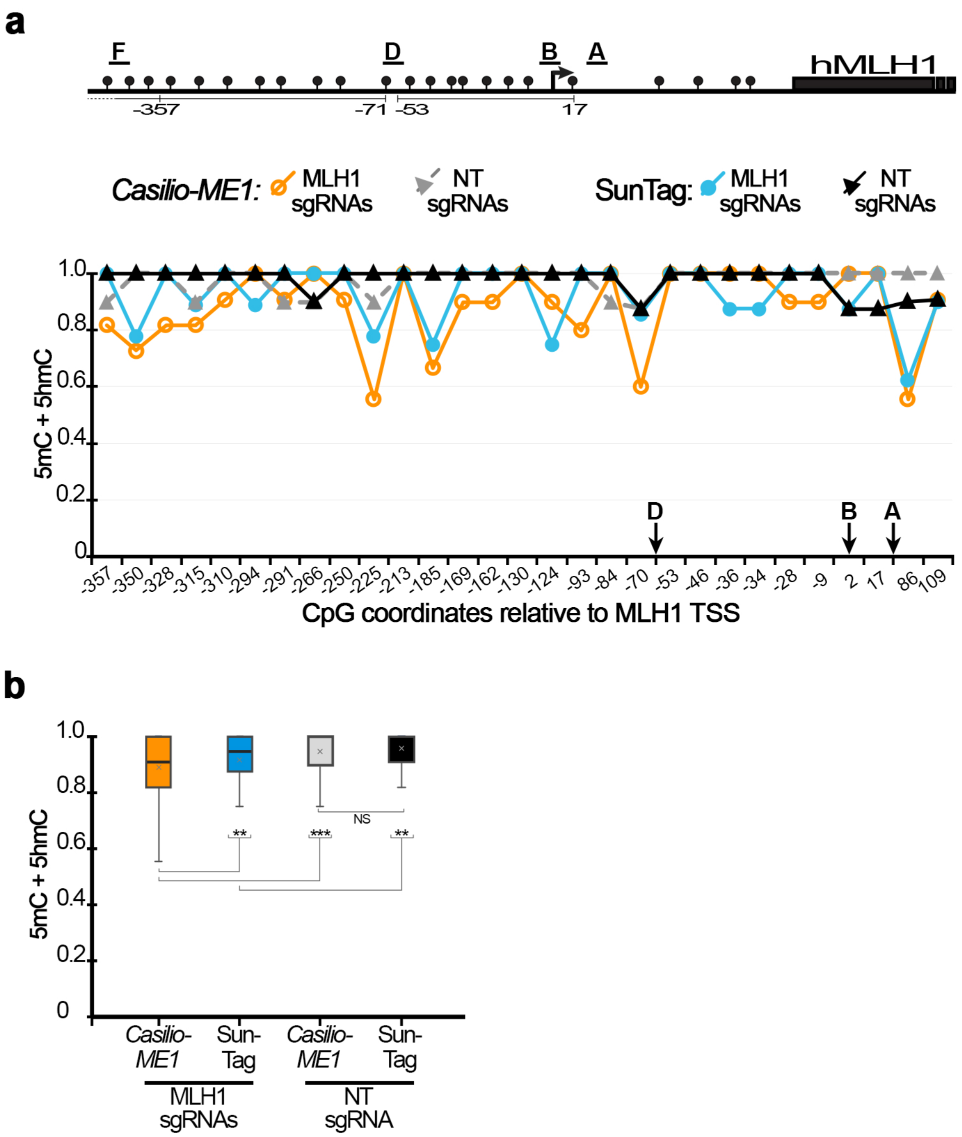

### Supplementary Figure 2

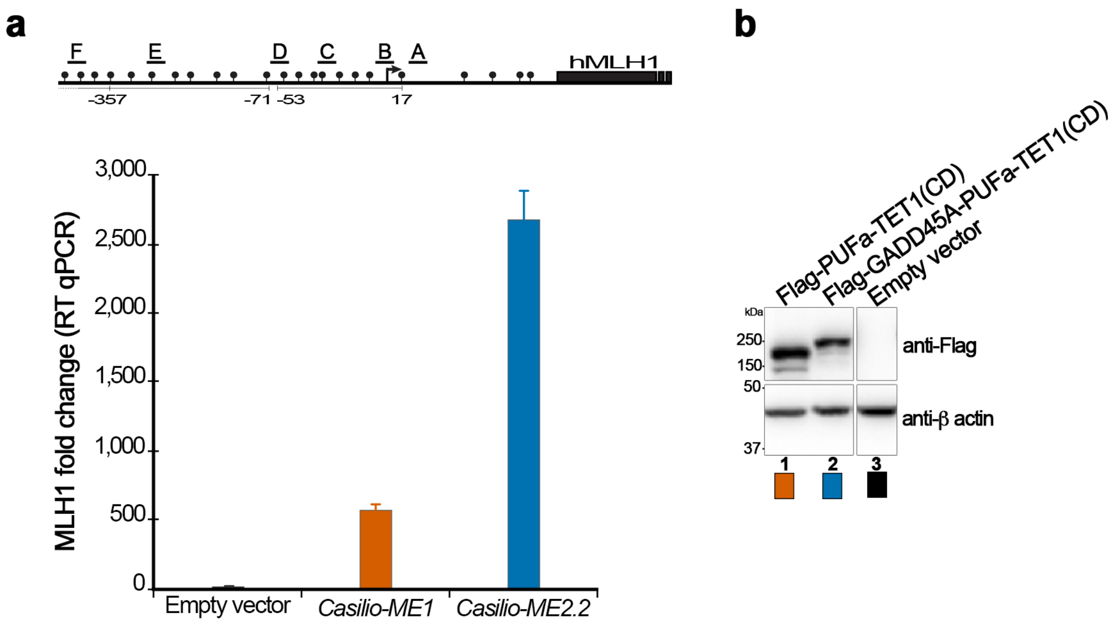

### Supplementary Figure 3

**a**

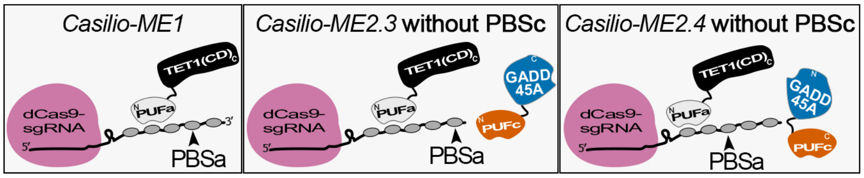


**b**

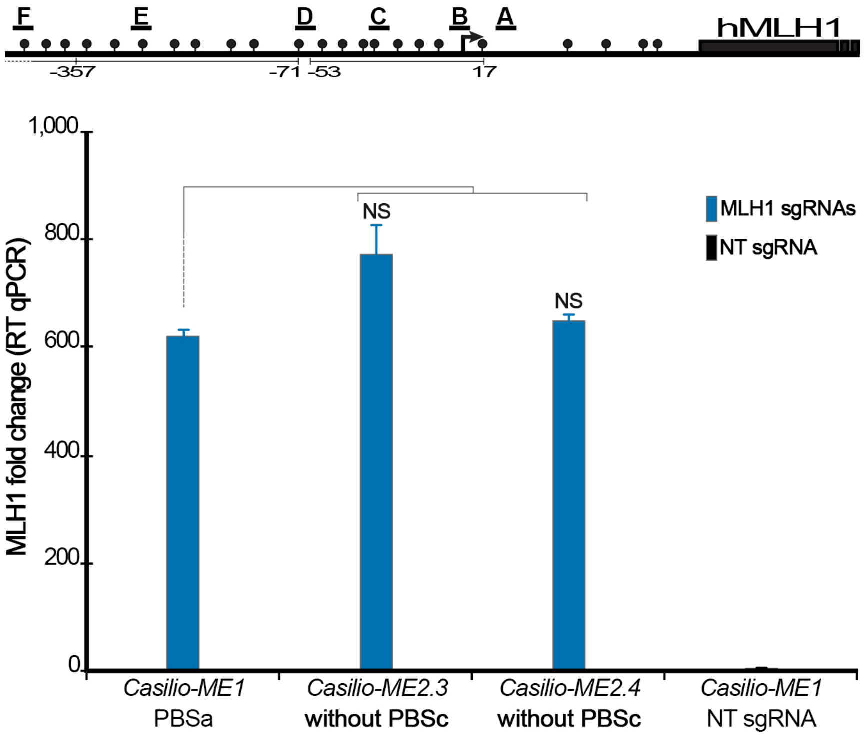

#### Supplementary Figure 4

**a**

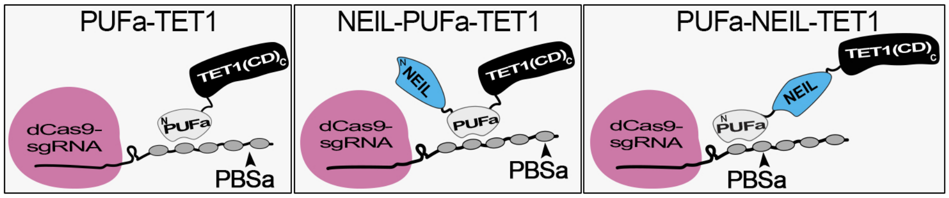


**b**

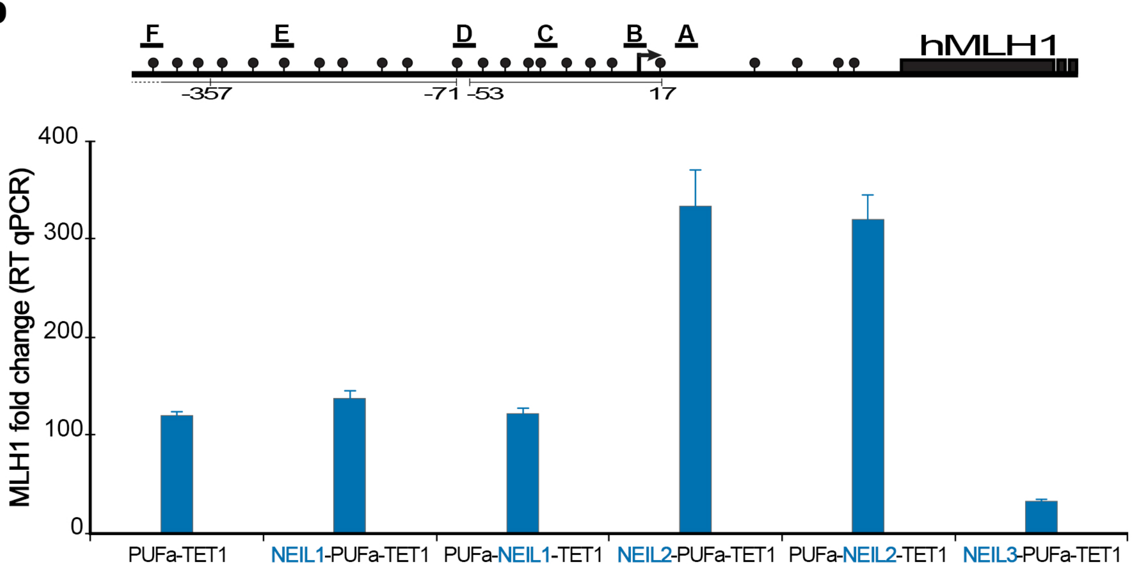


**c**

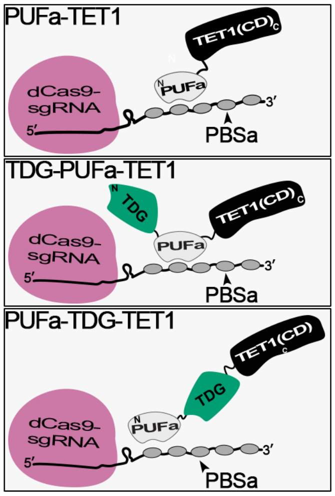


**d**

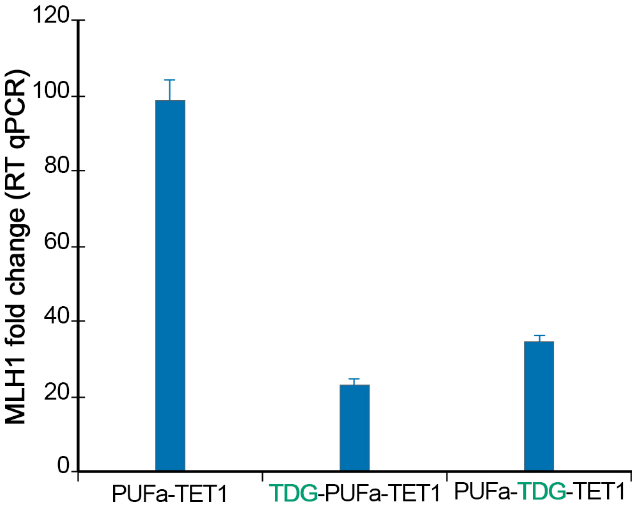

### Supplementary Figure 5

**a**

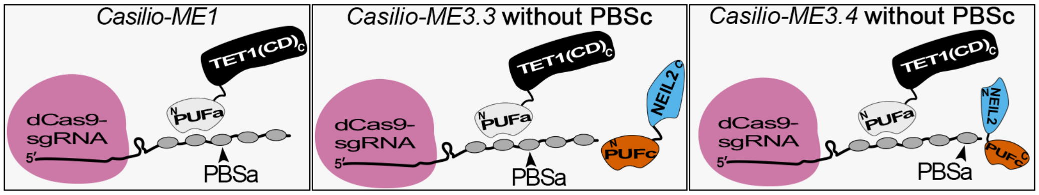


**b**

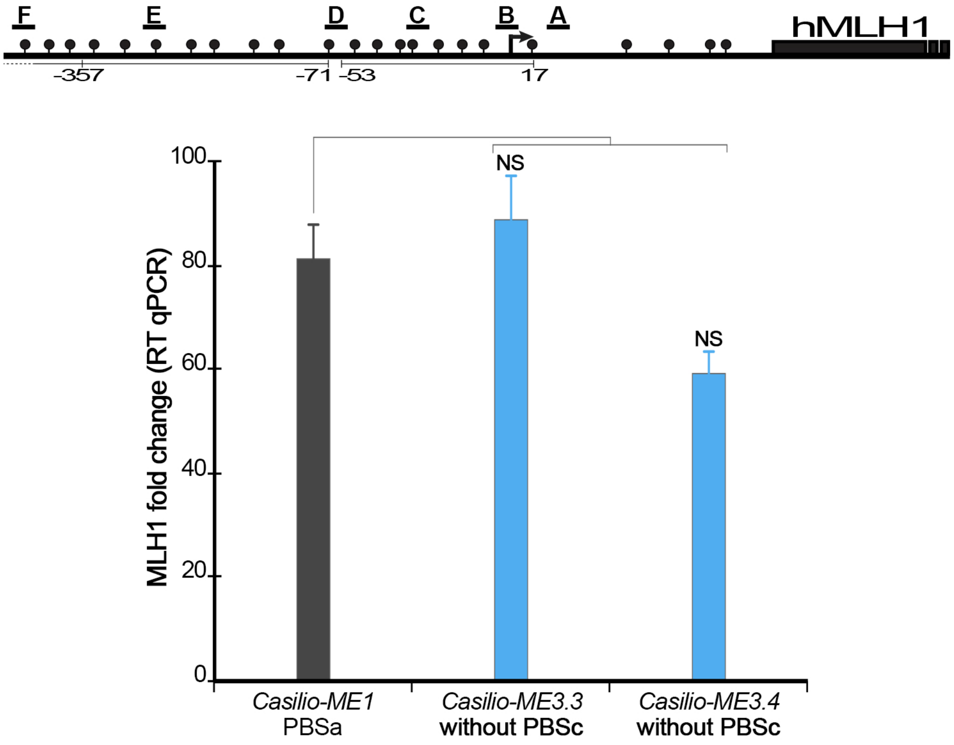

#### Supplementary Figure 6

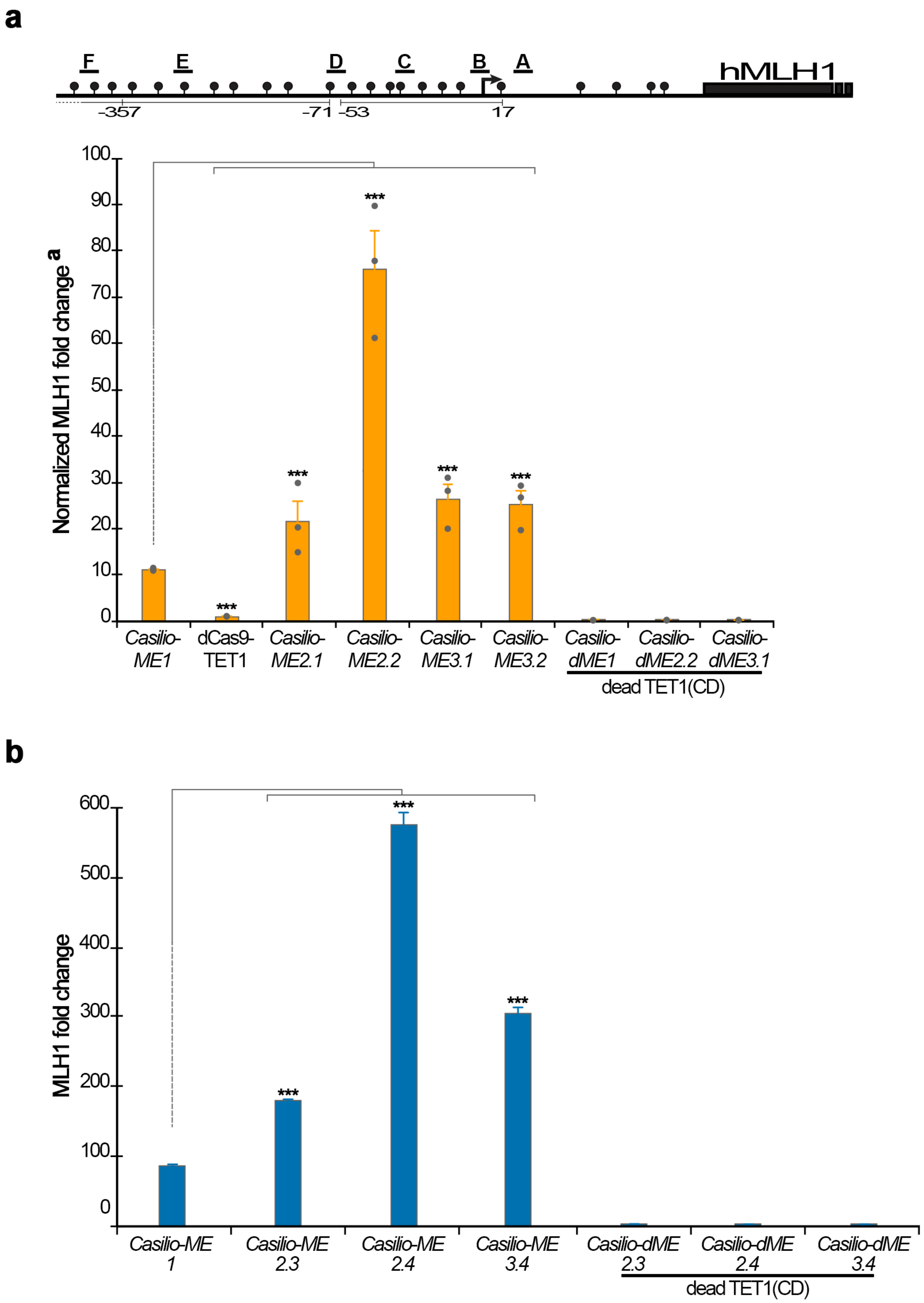

### Supplementary Figure 7

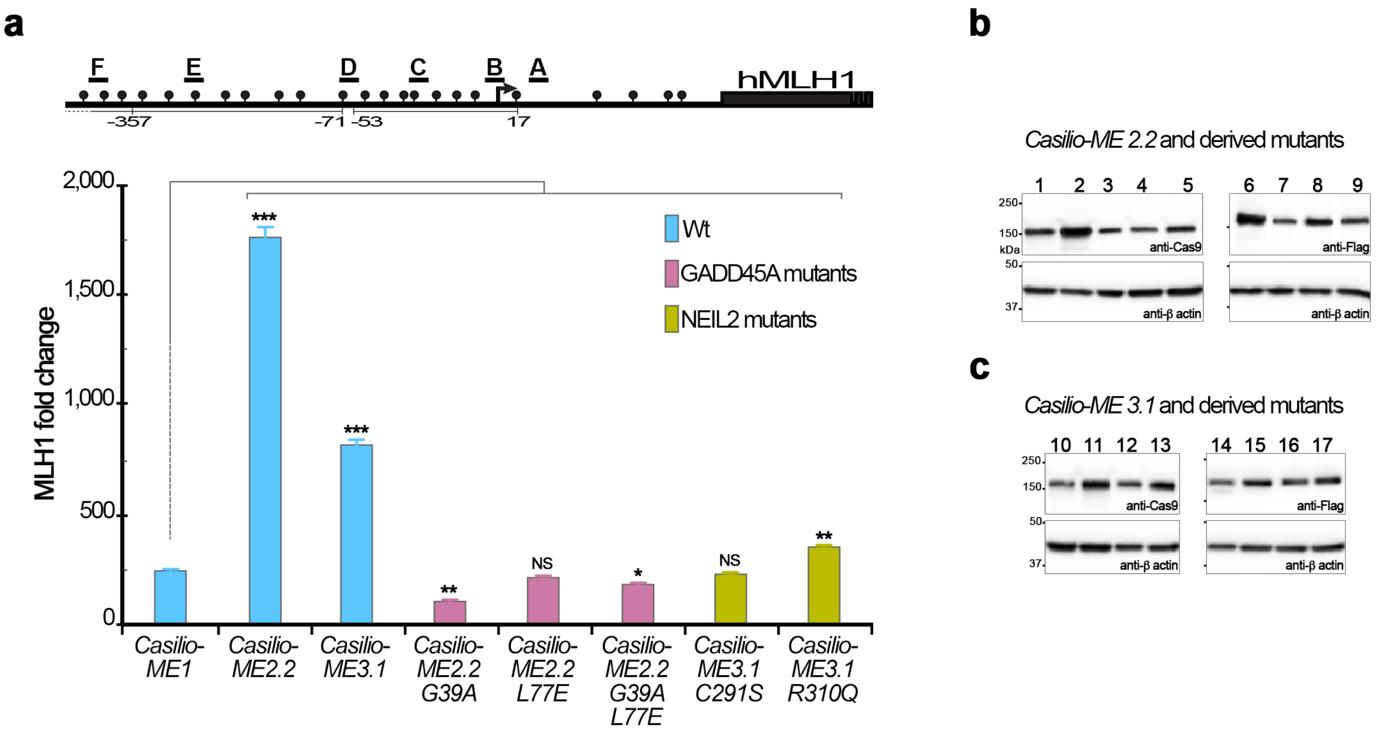

#### Supplementary Figure 8

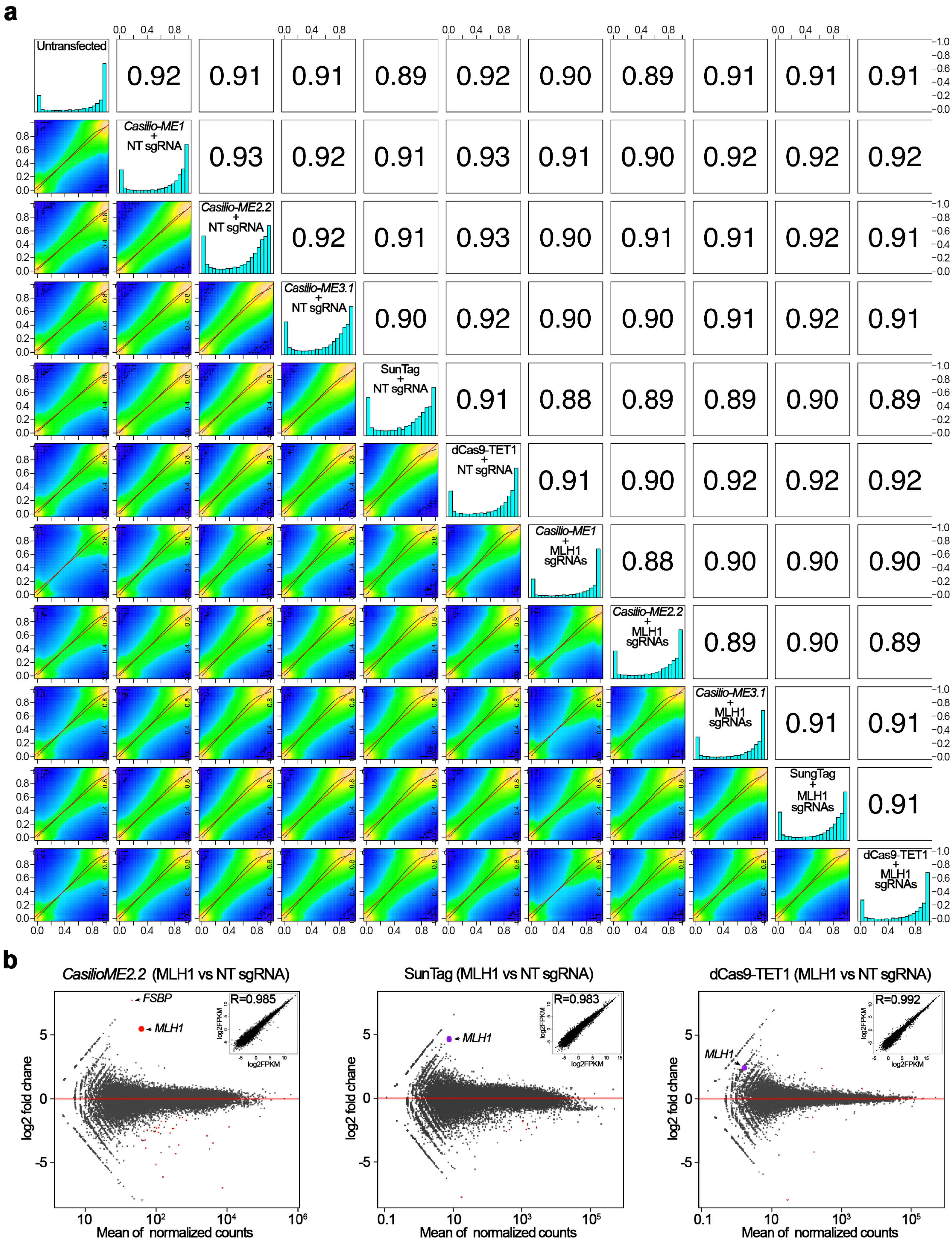

#### Supplementary Figure 9

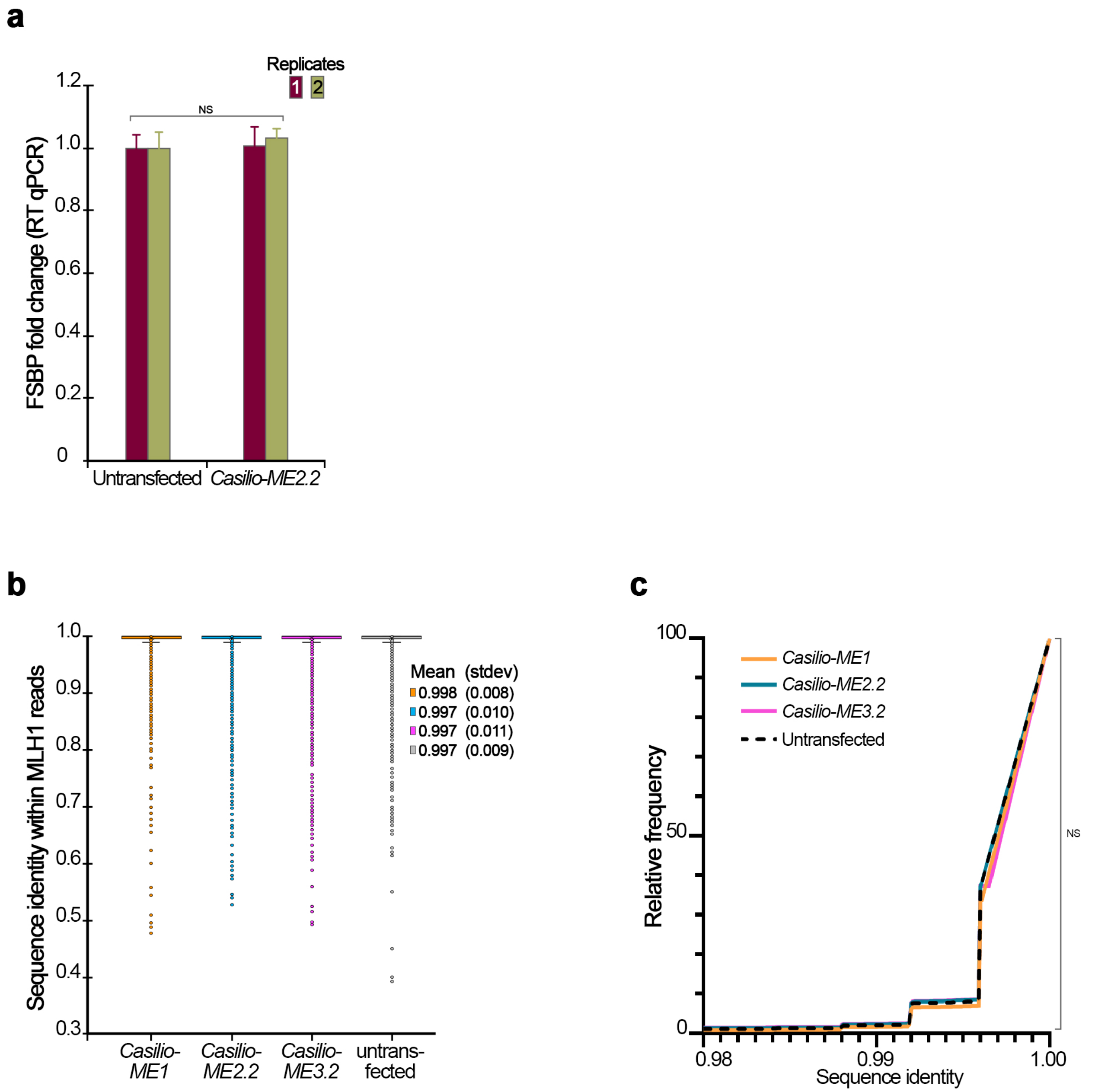

### Supplementary Figure 10

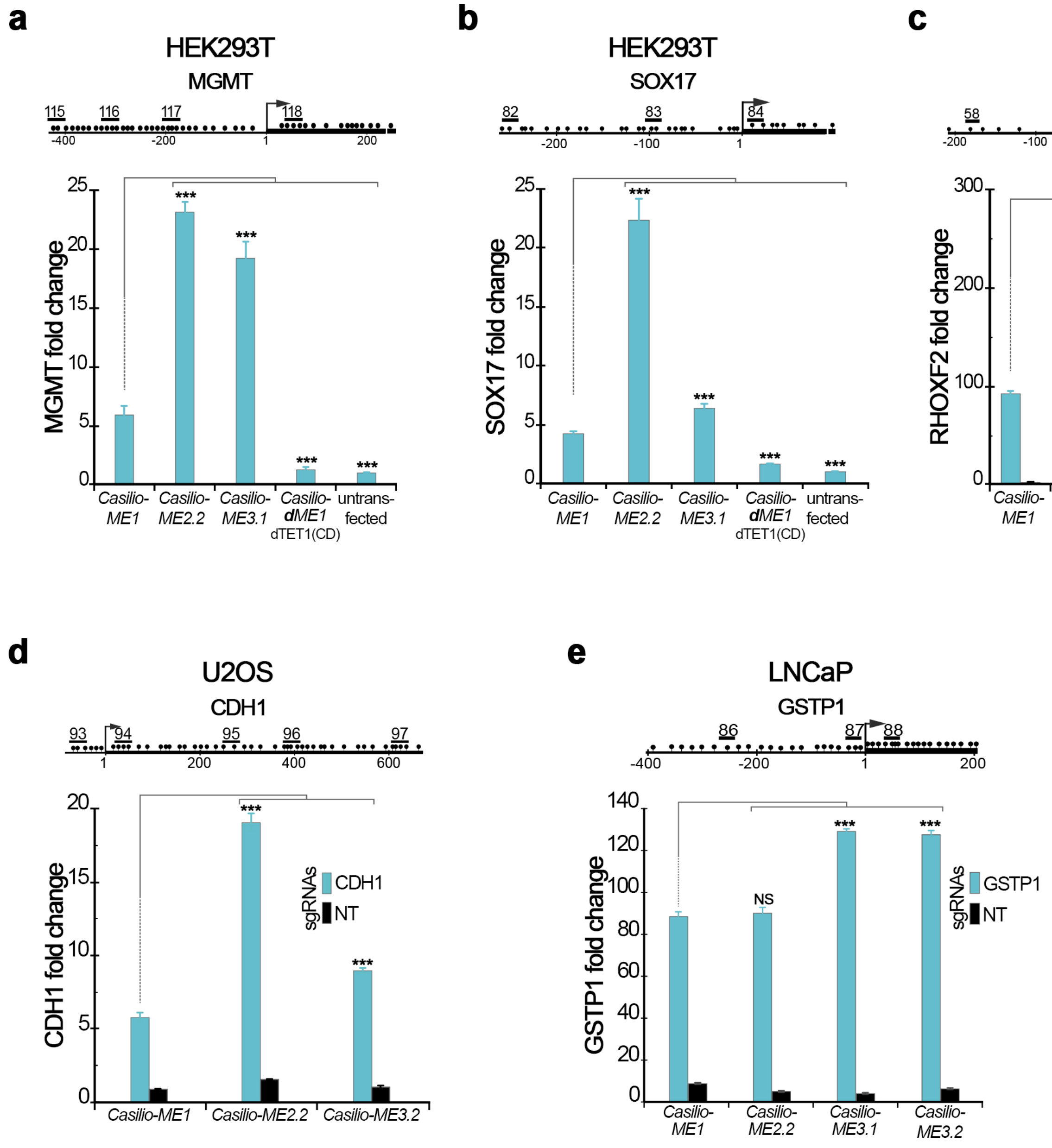

### Supplementary Figure 11

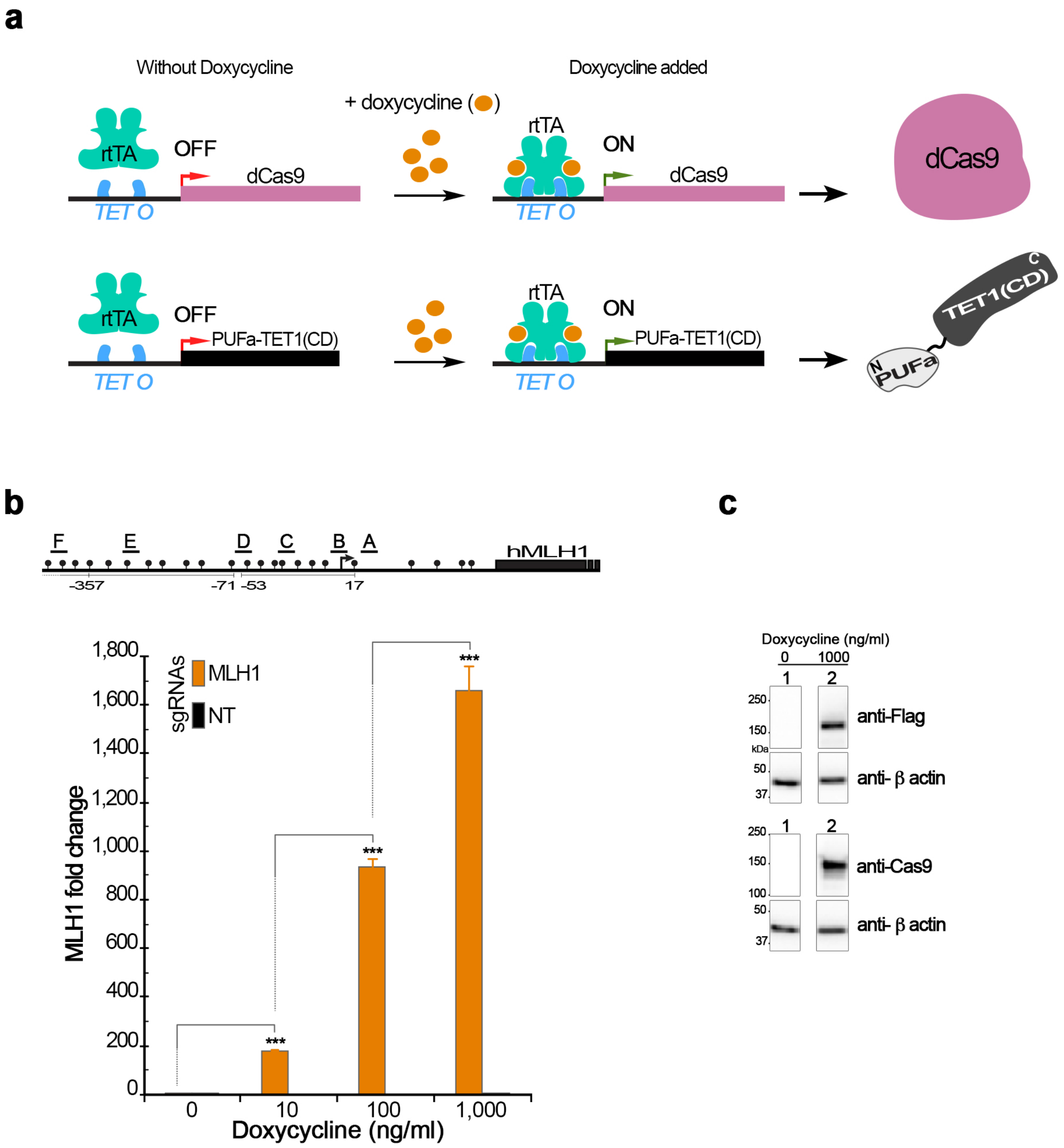

### SUPPLEMENTAL INFORMATION

#### LEGEND TO SUPPLEMENTARY FIGURES

**Supplementary Figure 1-** *Casilio-ME1* and SunTag mediated 5mC demethylation at *MLH1* promoter.

**(a)** Upper panel: *MLH1* promoter and associated CGI. CpGs (lollipops), TSS (arrow), and the sgRNAs used for targeting TET1 effectors (A, B, D, F) are shown.

Lower panel: methylation frequency (5mC + 5hmC) at *MLH1* promoter regions of cells transfected with components of *Casilio-ME1* or SunTag in the presence of *MLH1*-sgRNAs or NT-sgRNA is shown. Methylation frequency from BSeq of cloned *MLH1* amplicons plotted against CpG relative positions, and CpGs that overlap *MLH1*-sgRNAs sequence targets (arrows) are shown. Statistical significance of difference in methylation frequencies obtained were tested. $P<0.05$, one-way ANOVA.

**(b)** Box-plot of methylation frequency from BSeq of cloned *MLH1* amplicons from transfected cells shown in (a). NS, not significant, $P>0.05$, ** P<0.05, and *** P<0.001, one-way ANOVA.

**Supplementary Figure 2-** TET1-effector protein levels in cells transfected with *Casilio-ME1* or *Casilio-ME2.2* components showing that enhanced gene activation obtained by *Casilio-ME2.2* does not result from GADD45A-PUFa-TET1(CD) overexpression.

**(a)** *MLH1* mRNA relative levels (mean fold change ± S.E.M.; n=3) in cells transfected with components of Flag-tagged *Casilio-ME1* (Flag-PUFa-TET1(CD)) or

*Casilio-ME2.2* (Flag-GADD45A-PUFa-TET1(CD)) in the presence of indicated *MLH1*-sgRNAs depicted above the plot.

**(b)** Western blot analysis using anti-Flag, anti-β actin monoclonal antibodies and proteins extracted from cells analyzed (a). Proteins extracted from cells transfected with Flag-PUFa-TET1(CD) *Casilio-ME1* effector (lane 1), Flag-GADD45A-PUFa-TET1(CD) *Casilio-ME2.2* effector (lane 2), or empty vector (lane 3), and size of protein ladder in kDa are shown.

**Supplementary Figure 3-** Enhanced *MLH1* activation by *Casilio-ME2* platforms requires co-delivery of TET1(CD) and GADD45A effector modules to genomic sites.

**(a)** Illustration of *Casilio-ME* platforms showing effector modules of PUFa and PUFc protein fusions used to transfect cells in the presence of NT-sgRNA or *MLH1*-sgRNAs containing PBSa but lacking PBSc required for targeting PUFc-based effectors to genomic sites. TET1(CD) (black), GADD45A (blue), PUFa (light grey), PUFc (orange), amino (N) and carboxyl (C) termini of protein fusions are arbitrarily shown.

**(b)** *MLH1* mRNA relative levels (mean fold change ± S.E.M.; n=3) in cells transfected with the indicated components of *Casilio-ME1, Casilio-ME2.3, Casilio-ME2.4* in the presence of sgRNAs that comprised PBSa but lacked PBSc required for targeting PUFc-based effectors to target sites. The sgRNAs used to target *MLH1* promoter regions (A-F), CpGs (lollipops), and TSS (arrow) are depicted above the plot. NS, not significant, $P>0.05$, one-way ANOVA.

**Supplementary Figure 4-** Co-targeting TET1 activity and DNA glycosylases to *MLH1* promoter regions.

**(a)** Schematic representation of *Casilio-ME1* and its derivatives that include NEIL proteins (NEIL1, NEIL2 or NEIL3 glycosylases) as part of associated TET1 effector protein fusions. TET1(CD) (black), PUFa (light grey), NEIL proteins (blue), amino (N) and carboxyl (C) termini of protein fusion are arbitrarily shown.

**(b)** *MLH1* mRNA relative levels (mean fold change ± S.E.M.; n=3) as determined by TaqMan assays in cells transfected in the presence of indicated effectors. A drawing of sgRNAs used to target the *MLH1* promoter regions (A-F), CpGs (lollipops), and TSS (arrow) are shown above the plot. Similar results were obtained when NEIL1 and NEIL3 were used as modular PUFc effectors in the presence of PUFa-TET1(CD) and sgRNAs containing PBSa and PBSc.

**(c)** *Casilio-ME1* and its derivatives that include TDG as part of TET1 effector fusions are arbitrarily depicted to show TET1(CD) (black), PUFa (light grey), TDG (green), and amino (N) and carboxyl (C) termini of protein fusions.

**(d)** *MLH1* mRNA relative levels (mean fold change ± S.E.M.; n=3) in cells transfected in the presence of indicated effectors and the *MLH1*-sgRNAs shown in (b upper panel)**.** TDG failed to enhance *Casilio-ME* mediated gene activation when linked to TET1 effector although it plays an important role processing 5fC and 5caC. Same results were obtained when TDG was linked to PUFc as modular effectors in the presence of PUFa-TET1(CD) and sgRNAs containing PBSa and PBSc. This failure to enhance TET1-mediated gene activation with TDG could be due to improper folding of the TDG fusion proteins tested.

While alternative explanations exist, other factors might be required to permit coupling of TET1 and TDG activities at targeted sites.

**Supplementary Figure 5-** Enhanced *MLH1* activation by *Casilio-ME3* platforms requires co-delivery of TET1(CD) and NEIL2 effector modules to targeted genomic loci.

**(a)** Representation of *Casilio-ME* platforms showing effector modules of PUFa and PUFc protein fusions utilized to transfect cells in the presence of sgRNAs that contained PBSa but lacked PBSc required for targeting the PUFc-based effectors to genomic sites. TET1(CD) (black), NEIL2 (blue), PUFa (light grey), PUFc (orange), amino (N) and carboxyl (C) termini of protein fusions are arbitrarily shown.

**(b)** *MLH1* mRNA relative levels (mean fold change ± S.E.M.; n=3) in cells transfected with components of *Casilio-ME1, Casilio-ME3.3, Casilio-ME3.4* in the presence of sgRNAs that contained PBSa but lacked PBSc required for targeting the PUFc-based effectors to target sites. Drawings of the sgRNAs used to target the *MLH1* promoter regions (A-F), CpGs (lollipops), and TSS (arrow) are shown above the column plot. NS, not significant, $P>0.05$, one-way ANOVA.

**Supplementary Figure 6-** Comparison of the enhanced gene activation of *Casilio-ME2* and *Casilio-ME3* platforms.

**(a)** *MLH1* mRNA normalized levels (mean fold change ± S.E.M.; n=3) in cells transfected with components of *Casilio-ME1, Casilio-ME2.1, Casilio-ME2.2, Casilio-ME3.1, Casilio-ME3.2* or dCas9-TET1 systems in the presence of *MLH1*-sgRNAs. In the shown *Casilio-dME* derivatives, TET1(CD) was replaced by dTET1(CD) containing TET1-inactivating mutations. Drawings of the sgRNAs used to target the *MLH1* promoter regions (A-F), CpGs (lollipops), and TSS (arrow) are shown above the column plot. *** *P*< 0.0005, one-way ANOVA.

[a/] Fold changes were normalized to those obtained with targeting dCas9-TET1 to *MLH1* promoter which were 19.8 ± 0.7, 27.2 ± 1.8, and 16.7 ± 0.4 (mean fold change ± S.E.M.; n=3) relative to mock transfected cells in the three independent experiments shown, respectively.

**(b)** Relative levels *MLH1* mRNA (mean ± S.E.M.; n=3) in cells transfected with components of *Casilio-ME1, Casilio-ME2.3, Casilio-ME2.4* or *Casilio-ME3.4* as indicated. *MLH1*-sgRNA used are as shown in (a upper panel) but contained both PBSa and PBSc required for modular targeting of the associated effectors. dead TET1(CD) containing TET1-inactivating mutations replaced TET1(CD) in the shown *Casilio-dME* derivatives. *** *P*<0.0001, one-way ANOVA.

**Supplementary Figure 7-** Enhanced gene activations mediated by *Casilio-ME2* and *Casilio-ME3* platforms require active GADD45A and NEIL2 components.

**(a)** Column plot showing *MLH1* mRNA fold change (mean fold change ± S.E.M.; n=3) in cells transfected with components of *Casilio-ME1, Casilio-ME2.2, or Casilio-ME3.1* in the presence of the indicated *MLH1*-sgRNAs. Drawing of promoter regions with the *MLH1*-sgRNAs used (A-F), CpGs (lollipops), and TSS (arrow) is shown above the plot. Wild type (Wt) *Casilio-ME* platforms (blue) and derived *Casilio-ME2.2* and *Casilio-ME3.1* mutants containing the indicated point mutations that alter key functional properties of GADD45A (reddish purple) or inactive NEIL2 (green) are shown. NS, not significant, *P*>0.05, * *P*< 0.05, ** *P*<0.01, ****P*<0.0001, one-way ANOVA.

**(b, c)** Western blot analysis using anti-Cas9, anti-Flag or anti-β actin antibodies as indicated and protein extracts from cells analyzed for *MLH1* activation in (a). Flag-tagged and dCas9 protein components of wild type and derived mutants of *Casilio-ME* platforms *Casilio-ME2.2* (b) (lanes 1-9) and *Casilio-ME3.1* (c) (lanes 10-15) are shown: Wt (lanes 1, 6, 10 and 14), **d**TET1(CD) (lanes 2, 11 and 15), GADD45A(G39A) (lanes 3 and 7), GADD45A(L77E) (lanes 4 and 8), GADD45A(G39A L77E) (lanes 5 and 9), NEIL2(C291S) (lanes 12 and 16), and NEIL2(R310Q) (lanes 13 and 17).

**Supplementary Figure 8-** Evaluation of potential off-target effects on 5mC demethylation and gene expression.

**(a)** Histograms of CpG genome-wide 5mC methylation frequency and correlations between indicated samples with corresponding Pearson's correlation coefficient are shown. Untransfected cells or cells transfected with indicated 5mC demethylation systems in the presence of targeting *MLH1*-sgRNAs (A-F) or NT-sgRNA are shown.

**(b)** RNAseq analysis of two biological replicates of RNA samples obtained from cells transfected with *Casilio-ME2.2*, SunTag or dCas9-TET1 components in the presence of *MLH1*-sgRNAs or a NT-sgRNA are shown. Red dots represent differentially expressed hits deemed significant and purple dots represent non-significant *MLH1* differential expression based on obtained *P* values (see Methods). Pearson's correlation coefficients are shown as insert with corresponding MA-plots.

**Supplementary Figure 9-** Evaluation of potential off-target activation of *FSBP* expression and mutagenicity of *Casilio-ME* platforms.

**(a)** TaqMan assay showing *FSBP* mRNA relative levels (mean fold change ± S.E.M; n=3) in the two replicates of mRNA samples analyzed by RNAseq shown in (Fig. S8b). NS, not significant, $P>0.05$, two-way ANOVA.

**(b**, **c)** Sequence analysis of *MLH1* reads in amplicons obtained from untransfected cells or cells transfected with components of the indicated *Casilio-ME* platforms and the six *MLH1*-sgRNAs. Box plot (b) and cumulative distribution frequency plot (c) of sequence identity distribution in *MLH1* reads. Statistical significance of differences in sequence identity distribution between samples were tested. NS, not significant, $P>0.5$, one-way ANOVA. $P>0.1$ (untransfected cells vs *Casilio-ME1*), $P>0.5$ (untransfected cells vs *Casilio-ME2.2*), and $P>0.1$ (untransfected cells vs *Casilio-ME3.2*), Mann-Whitney U test.

**Supplementary Figure 10-** *Casilio-ME* platforms enable enhanced activation of methylation-regulated genes.

**(a-e)** mRNA fold changes (mean fold change ± S.E.M.; n=3) obtained by targeting different genes in different cell types by transfection (HEK293T, U2OS) or nucleofection (LNCaP) of indicated *Casilio-ME* components and sgRNAs are shown. mRNA levels were quantitated in cells collected 3 days after transfection. When indicated TET1(CD) was replaced with catalytically dead TET(CD) to show the role of TET1 oxidative activity in the obtained gene activations. The targeted promoter regions with associated CGI depicting TSS (arrow), CpG (lollipops) and location of sgRNAs used (lines under numbers) are shown above corresponding plot. NS, not significant, $P > 0.05$, *** $P < 0.005$, one-way ANOVA.

**Supplementary Figure 11-** Tunable activation of methylation-silenced genes.

**(a)** Depiction of expression inducible *Casilio-ME1* components using Tet-ON system. The reverse tetracycline-controlled transcriptional activator (r-tTA) does not bind *Tet* operator (TET O) sequences in the absence of doxycycline. Doxycycline-bound rtTA undergoes conformational changes allowing its binding to TET O sequences and inducing subsequent transcriptional activation of associated genes.

**(b)** Range of *MLH1* activation (mean mRNA fold change ± S.E.M.; n=3) obtained with *DIP_Casilio-ME1* platform when media were supplemented with a range of Dox concentration. PiggyBac vectors hosting cassette enabling a Dox-inducible expression of dCas9 or PUFa-TET1(CD) effector were concomitantly delivered to cells via PiggyBac transposase system[1 2], and doubly selected cells were then transiently transfected with *MLH1*-sgRNAs. Depiction of *MLH1* promoter showing sgRNAs used is shown above the plot. *** $P<0.0001$, one-way ANOVA.

**(c)** Western blot analysis using anti-Flag, anti-Cas9, anti-β actin monoclonal antibodies and protein extracts form DIP_*Casilio-ME1* cells transfected with *MLH1*-sgRNAs without (lanes 1) or with 1μg/ml (lane 2) supplemented doxycycline is shown.

**TABLE 1-** Plasmid list

| Plasmid ID | Description | Addgene ID |
|---|---|---|
| pAC1371 | pX-sgRNA-5xPBSa- Cloning vector to express sgRNA-5xPBSa | 71888 |
| pAT243 | pX-sgRNA-5xPBSa- Cloning vector to express sgRNA-5xPBSa_5xPBSc | Pending |
| pAT888 | CMV/CAG_PUFa-hTET1(CD) | Pending |
| pAC1445 | pmax_dCas9 | 73169 |
| pAT890 | CMV/CAG_dCas9-hTET1(CD) | Pending |
| pCAG-dCas9-5xPlat2AflD | For dCas9-(SunTag)x5 array expression | 82560 |
| pCAG-scFvGCN4sfGFPTET1CD | For antibody-sfGFP-hTET1(CD) expression | 82561 |
| pcDNA3.1-MS2-Tet1-CD | For MS2 coat protein-mTET1(CD) expression | 83341 |
| pdCas9-Tet1-CD | For dCas9-TET1(CD) and sgRNA expression | 83340 |
| pAT801 | CMV/CAG_TALE_A-hTET1(CD) | Pending |
| pAT812 | CMV/CAG_TALE_B-hTET1(CD) | Pending |
| pAT806 | CMV/CAG_TALE_D-hTET1(CD) | Pending |
| pAT817 | CMV/CAG_TALE_F-hTET1(CD) | Pending |
| pAT341 | CMV/CAG_PUFa-TET1CD (H1671Y D1673A) (dead TET1) | Pending |
| pAT360 | CMV/CAG_PUFa-hGADD45A-hTET1(CD) | Pending |
| pAT635 | CMV/CAG_hGADD45A-PUFa-hTET1(CD) | Pending |
| pAT355 | CMV/CAG_PUFc-hGADD45A | Pending |
| pAT356 | CMV/CAG_hGADD45A-PUFc | Pending |
| pAT892 | CMV/CAG_PUFa-p65HSF1 | Pending |
| pAT608 | CMV/CAG_Flag-hNEIL2-PUFa-hTET1(CD) | Pending |
| pAT611 | CMV/CAG_PUFa-hNEIL2-hTET1(CD) | Pending |
| pAT595 | CMV/CAG_PUFc-hNEIL2 | Pending |
| pAT596 | CMV/CAG_hNEIL2-PUFc | Pending |
| pAT607 | CMV/CAG_hNEIL1-PUFa-hTET1(CD) | N/A |
| pAT610 | CMV/CAG_PUFa-hNEIL1-hTET1(CD) | N/A |
| pAT609 | CMV/CAG_hNEIL3-PUFa-hTET1(CD) | N/A |
| pAT359 | CMV/CAG_PUFa-hTDG-hTET1(CD) | N/A |

| | | |
|---|---|---|
| pAT335 | CMV/CAG_hTDG-PUFa-hTET1(CD) | N/A |
| pAT699 | CMV/CAG_Flag-hGADD45A-PUFa-hTET1(CD) | Pending |
| pAT976 | CMV/CAG_Flag-hGADD45A-PUFa-dTET1CD | Pending |
| pAT971 | CMV/CAG_Flag-hGADD45A (G39A)-PUFa-hTET1(CD**)** | Pending |
| pAT972 | CMV/CAG_Flag-hGADD45A (L77E)-PUFa-hTET1(CD**)** | Pending |
| pAT973 | CMV/CAG_FlaghGADD45A (G39A, L77E)-PUFa-hTET1(CD**)** | Pending |
| pAT977 | CMV/CAG_Flag-hNEIL2-PUFa-dTET1(CD) | Pending |
| pAT1059 | CMV/CAG_Flag-hNEIL2 (C291S)-PUFa-hTET1(CD) | Pending |
| pAT1060 | CMV/CAG_Flag-hNEIL2 (R310Q)-PUFa-hTET1(CD) | Pending |
| pAT1089 | PB-EF1a-Blast_2A_rtTA3-SV40pA_pCW-dCas9 | Pending |
| pAT1090 | PB-EF1a-Hygro_2A_rtTA3-SV40pA_pCW-Flag-PUFa-hTET1(CD) | Pending |

**TABLE 2-** List of sgRNA spacer sequences.

| sgRNAs | spacer sequences | References |
|---|---|---|
| MLH1-A | gACAGAGTTGAGAAATTTGAC | This work |
| MLH1-B | GGCAGTAGCCGCTTCAGGGA | This work |
| MLH1-C | GCGCAAGCGCATATCCTTCT | This work |
| MLH1-D | gAAACGAACCAATAGGAAGAG | This work |
| MLH1-E | GCGCCAGATCACCTCAGCAG | This work |
| MLH1-F | gCTGACGCAGACGCTCCACCA | This work |
| Non targeting | gTTCTCTTGCTGAAAGCTCGA | [3] |
| RHOXF2-57 | gCCCGCTATTTGCTGTGGGTT | [4] |
| RHOXF2-58 | gACTCACGCATGCCTGTCTAC | This work |
| RHOXF2-59 | gTAGCACTGCCTAGGAGAGCG | This work |
| RHOXF2-60 | gAACGCGTGCTCTCCCTCATC | This work |
| SOX17-82 | GTACAATCAGCCCTCCCAGA | This work |
| SOX17-83 | GTGGGACTCGGACCACGGCC | This work |
| SOX17-84 | gTCTGTGCAGAAAAGGCCCCG | This work |
| GSTP1-85 | GAAGCGGGTGTGCAAGCTCC | This work |
| GSTP1-86 | GTTTACTCCCTAGGCCCCGC | This work |
| GSTP1-87 | gTATAAGGCTCGGAGGCCGCG | This work |
| GSTP1-88 | gTCGCCACCAGTGAGTACGCG | This work |
| CDH1-93 | GCGTCTATGCGAGGCCGGGT | This work |
| CDH1-94 | GTACGGGGGGCGGTGCCTCCG | This work |
| CDH1-95 | gCCGGATCCCCTGACTTGCGA | This work |
| CDH1-96 | GCCTGGAAGCCTCGCGCGCTC | This work |
| CDH1-97 | gAGTCGTGGGGACGATCTTCG | This work |
| MGMT-114 | GGACCGGGATTCTCACTAAG | This work |
| MGMT-115 | GCAGGTCGCTTGCACGCCCG | This work |
| MGMT-116 | GCCCGGCTTGTACCGGCCGA | This work |
| MGMT-117 | GCACAGGGCATGCGCCGACC | This work |

**TABLE 3-** Sequence of the sgRNA scaffold with 5xPBSa and 5xPBSc

| Description | DNA Sequence |
|---|---|
| U6 promoter-sgRNA-5x**PBSa-**PBSc For expression of modified sgRNAs comprising spacer sequence (*N*s) inserted at the 5'end, and a set of five binding sites for PUFa (PBSa, bold face) or PUFc (PBSc, underlined) added to the 3'region of the sgRNA. | gagggcctatttcccatgattccttcatatttgcatatacgatacaaggctgttagagagataattggaattaatttgactgtaaacacaaagatattagtacaaaatacgtgacgtagaaagtaataatttcttgggtagtttgcagttttaaaattatgttttaaaatggactatcatatgcttaccgtaacttgaaagtatttcgatttcttggctttatatatcttGTGGAAAGGACGAAACACC*gNNNNNNNNNNNNNNNNNNN*GTTTaagagctaTGCTGGAAACAGCAtagcaagttTaaataaggctagtccgttatcaacttgaaaaagtggcaccgagtcggtgcCAATTGggtctccagat**TGTATGTA**gcc**TGTATGTA**gcc**TGTATGTA**gcc**TGTATGTA**gcc**TGTATGTA**aGATCCAATTGggtctccagatTTGATGTAgccTTGATGTAgccTTGATGTAgccTTGATGTAgccTTGATGTAagatCTTTTTTTgttttagagctagaaatagcaagttaaaataaggctagtccgtagcgcgtgcgccaattctgcagacaaatggc |

U6-sgRNA containing 5 PBSa, Addgene ID #71888 was as previously reported [3].

**TABLE 4-** List of bisulfite sequencing PCR primer

| Primers | Primer sequences | Targeted promoters |
|---|---|---|
| AT442-F | gatccccgggtaccgagctcgaattAAGGTTAAGAGGYGGTAGAGTT | *MLH1* distal (cloning) |
| AT443-R | cgttgtaaaacgacggccagtgaattTTAACCCTACTCTTATAACCTCCC | |
| AT444-F | gatccccgggtaccgagctcgaattGGGAGGTTATAAGAGTAGGGTTAA | *MLH1* intermediate (cloning) |
| AT445-R | cgttgtaaaacgacggccagtgaattCATCCAACCCCACCCTTCAA | |
| AT446-F | gatccccgggtaccgagctcgaattTTGAAGGGTGGGGTTGGATG | *MLH1* proximal (cloning) |
| AT447-R | cgttgtaaaacgacggccagtgaattTTATAAACATACRCTATACATACCTCTACC | |
| AT531-F | AAGGTTAAGAGGYGGTAGAGTT | *MLH1* distal |
| AT532-R | TTAACCCTACTCTTATAACCTCCC | |
| AT533-F | GGGAGGTTATAAGAGTAGGGTTAA | *MLH1* intermediate |
| AT534-R | CATCCAACCCCACCCTTCAA | |
| AT535-F | TTGAAGGGTGGGGTTGGATG | *MLH1* proximal |
| AT536-R | TTATAAACATACRCTATACATACCTCTACC | |

**TABLE 5-** List of relevant protein sequences

| |
|---|
| Name: **PUFa-TET1(CD)** |
| **Keys:** NLS, PUFa, **TET1(1418-2136)** |
| MIDGGGGSDPKKKRKVDPKKKRKVDPKKKRKVGSTGSRNDGGGGSGGGGSGGGGSGRAGILPPKKK<br>RKVSRGRSRLLEDFRNNRYPNLQLREIAGHIMEFSQDQHGSRFIQLKLERATPAERQLVFNEILQAAYQL<br>MVDVFGNYVIQKFFEFGSLEQKLALAERIRGHVLSLALQMYGSRVIEKALEFIPSDQQNEMVRELDGHVL<br>KCVKDQNGNHVVQKCIECVQPQSLQFIIDAFKGQVFALSTHPYGCRVIQRILEHCLPDQTLPILEELHQHT<br>EQLVQDQYGNYVIQHVLEHGRPEDKSKIVAEIRGNVLVLSQHKFASNVVEKCVTHASRTERAVLIDEVCT<br>MNDGPHSALYTMMKDQYANYVVQKMIDVAEPGQRKIVMHKIRPHIATLRKYTYGKHILAKLEKYYMKNGV<br>DLGDPKKKRKVDPKKKRKVGGRGGGGSGGGGSGGGGSGPA**ELPTCSCLDRVIQKDKGPYYTHLGAG**<br>**PSVAAVREIMENRYGQKGNAIRIEIVVYTGKEGKSSHGCPIAKWVLRRSSDEEKVLCLVRQRTGHHCPT**<br>**AVMVVLIMVWDGIPLPMADRLYTELTENLKSYNGHPTDRRCTLNENRTCTCQGIDPETCGASFSFGCS**<br>**WSMYFNGCKFGRSPSPRRFRIDPSSPLHEKNLEDNLQSLATRLAPIYKQYAPVAYQNQVEYENVAREC**<br>**RLGSKEGRPFSGVTACLDFCAHPHRDIHNMNNGSTVVCTLTREDNRSLGVIPQDEQLHVLPLYKLSDT**<br>**DEFGSKEGMEAKIKSGAIEVLAPRRKKRTCFTQPVPRSGKKRAAMMTEVLAHKIRAVEKKPIPRIKRKN**<br>**NSTTTNNSKPSSLPTLGSNTETVQPEVKSETEPHFILKSSDNTKTYSLMPSAPHPVKEASPGFSWSPKT**<br>**ASATPAPLKNDATASCGFSERSSTPHCTMPSGRLSGANAAAADGPGISQLGEVAPLPTLSAPVMEPLI**<br>**NSEPSTGVTEPLTPHQPNHQPSFLTSPQDLASSPMEEDEQHSEADEPPSDEPLSDDPLSPAEEKLPHI**<br>**DEYWSDSEHIFLDANIGGVAIAPAHGSVLIECARRELHATTPVEHPNRNHPTRLSLVFYQHKNLNKPQH**<br>**GFELNKIKFEAKEAKNKKMKASEQKDQAANEGPEQSSEVNELNQIPSHKALTLTHDNVVTVSPYALTH**<br>**VAGPYNHWV**ID |
| Name: **dCas9** |
| **Keys:** NLS, *Sp* dCas9, HA tag |
| MIDGGGGSGGGGSGGGGSMYPYDVPDYASPKKKRKVEASDKKYSIGLAIGTNSVGWAVITDEYKVPSK<br>KFKVLGNTDRHSIKKNLIGALLFDSGETAEATRLKRTARRRYTRRKNRICYLQEIFSNEMAKVDDSFFHRL<br>EESFLVEEDKKHERHPIFGNIVDEVAYHEKYPTIYHLRKKLVDSTDKADLRLIYLALAHMIKFRGHFLIEGDL<br>NPDNSDVDKLFIQLVQTYNQLFEENPINASGVDAKAILSARLSKSRRLENLIAQLPGEKKNGLFGNLIALSL<br>GLTPNFKSNFDLAEDAKLQLSKDTYDDDLDNLLAQIGDQYADLFLAAKNLSDAILLSDILRVNTEITKAPLSA<br>SMIKRYDEHHQDLTLLKALVRQQLPEKYKEIFFDQSKNGYAGYIDGGASQEEFYKFIKPILEKMDGTEELL<br>VKLNREDLLRKQRTFDNGSIPHQIHLGELHAILRRQEDFYPFLKDNREKIEKILTFRIPYYVGPLARGNSRF<br>AWMTRKSEETITPWNFEEVVDKGASAQSFIERMTNFDKNLPNEKVLPKHSLLYEYFTVYNELTKVKYVTE<br>GMRKPAFLSGEQKKAIVDLLFKTNRKVTVKQLKEDYFKKIECFDSVEISGVEDRFNASLGTYHDLLKIIKDK<br>DFLDNEENEDILEDIVLTLTLFEDREMIEERLKTYAHLFDDKVMKQLKRRRYTGWGRLSRKLINGIRDKQS<br>GKTILDFLKSDGFANRNFMQLIHDDSLTFKEDIQKAQVSGQGDSLHEHIANLAGSPAIKKGILQTVKVVDEL<br>VKVMGRHKPENIVIEMARENQTTQKGQKNSRERMKRIEEGIKELGSQILKEHPVENTQLQNEKLYLYYLQ<br>NGRDMYVDQELDINRLSDYDVDAIVPQSFLKDDSIDNKVLTRSDKNRGKSDNVPSEEVVKKMKNYWRQL<br>LNAKLITQRKFDNLTKAERGGLSELDKAGFIKRQLVETRQITKHVAQILDSRMNTKYDENDKLIREVKVITLK<br>SKLVSDFRKDFQFYKVREINNYHHAHDAYLNAVVGTALIKKYPKLESEFVYGDYKVYDVRKMIAKSEQEIG<br>KATAKYFFYSNIMNFFKTEITLANGEIRKRPLIETNGETGEIVWDKGRDFATVRKVLSMPQVNIVKKTEVQT<br>GGFSKESILPKRNSDKLIARKKDWDPKKYGGFDSPTVAYSVLVVAKVEKGKSKKLKSVKELLGITIMERSS<br>FEKNPIDFLEAKGYKEVKKDLIIKLPKYSLFELENGRKRMLASAGELQKGNELALPSKYVNFLYLASHYEKL<br>KGSPEDNEQKQLFVEQHKHYLDEIIEQISEFSKRVILADANLDKVLSAYNKHRDKPIREQAENIIHLFTLTNL<br>GAPAAFKYFDTTIDRKRYTSTKEVLDATLIHQSITGLYETRIDLSQLGGDSPKKKRKVEASGGGGSGGGG<br>SGGGGSGPA |
| **Name: dCas9-TET1(CD)** |
| **Keys:** NLS, *Sp* dCas9, **TET1(1418-2136),** HA tag |
| MIDGGGGSGGGGSGGGGSMYPYDVPDYASPKKKRKVEASDKKYSIGLAIGTNSVGWAVITDEYKVPSK<br>KFKVLGNTDRHSIKKNLIGALLFDSGETAEATRLKRTARRRYTRRKNRICYLQEIFSNEMAKVDDSFFHRL<br>EESFLVEEDKKHERHPIFGNIVDEVAYHEKYPTIYHLRKKLVDSTDKADLRLIYLALAHMIKFRGHFLIEGDL |

NPDNSDVDKLFIQLVQTYNQLFEENPINASGVDAKAILSARLSKSRRLENLIAQLPGEKKNGLFGNLIALSL
GLTPNFKSNFDLAEDAKLQLSKDTYDDDLDNLLAQIGDQYADLFLAAKNLSDAILLSDILRVNTEITKAPLSA
SMIKRYDEHHQDLTLLKALVRQQLPEKYKEIFFDQSKNGYAGYIDGGASQEEFYKFIKPILEKMDGTEELL
VKLNREDLLRKQRTFDNGSIPHQIHLGELHAILRRQEDFYPFLKDNREKIEKILTFRIPYYVGPLARGNSRF
AWMTRKSEETITPWNFEEVVDKGASAQSFIERMTNFDKNLPNEKVLPKHSLLYEYFTVYNELTKVKYVTE
GMRKPAFLSGEQKKAIVDLLFKTNRKVTVKQLKEDYFKKIECFDSVEISGVEDRFNASLGTYHDLLKIIKDK
DFLDNEENEDILEDIVLTLTLFEDREMIEERLKTYAHLFDDKVMKQLKRRRYTGWGRLSRKLINGIRDKQS
GKTILDFLKSDGFANRNFMQLIHDDSLTFKEDIQKAQVSGQGDSLHEHIANLAGSPAIKKGILQTVKVVDEL
VKVMGRHKPENIVIEMARENQTTQKGQKNSRERMKRIEEGIKELGSQILKEHPVENTQLQNEKLYLYYLQ
NGRDMYVDQELDINRLSDYDVDAIVPQSFLKDDSIDNKVLTRSDKNRGKSDNVPSEEVVKKMKNYWRQL
LNAKLITQRKFDNLTKAERGGLSELDKAGFIKRQLVETRQITKHVAQILDSRMNTKYDENDKLIREVKVITLK
SKLVSDFRKDFQFYKVREINNYHHAHDAYLNAVVGTALIKKYPKLESEFVYGDYKVYDVRKMIAKSEQEIG
KATAKYFFYSNIMNFFKTEITLANGEIRKRPLIETNGETGEIVWDKGRDFATVRKVLSMPQVNIVKKTEVQT
GGFSKESILPKRNSDKLIARKKDWDPKKYGGFDSPTVAYSVLVVAKVEKGKSKKLKSVKELLGITIMERSS
FEKNPIDFLEAKGYKEVKKDLIIKLPKYSLFELENGRKRMLASAGELQKGNELALPSKYVNFLYLASHYEKL
KGSPEDNEQKQLFVEQHKHYLDEIIEQISEFSKRVILADANLDKVLSAYNKHRDKPIREQAENIIHLFTLTNL
GAPAAFKYFDTTIDRKRYTSTKEVLDATLIHQSITGLYETRIDLSQLGGDSPKKKRKVEASGGGGSGGGG
SGGGGSGPA**ELPTCSCLDRVIQKDKGPYYTHLGAGPSVAAVREIMENRYGQKGNAIRIEIVVYTGKEGK**
**SSHGCPIAKWVLRRSSDEEKVLCLVRQRTGHHCPTAVMVVLIMVWDGIPLPMADRLYTELTENLKSYN**
**GHPTDRRCTLNENRTCTCQGIDPETCGASFSFGCSWSMYFNGCKFGRSPSPRRFRIDPSSPLHEKNLE**
**DNLQSLATRLAPIYKQYAPVAYQNQVEYENVARECRLGSKEGRPFSGVTACLDFCAHPHRDIHNMNNG**
**STVVCTLTREDNRSLGVIPQDEQLHVLPLYKLSDTDEFGSKEGMEAKIKSGAIEVLAPRRKKRTCFTQP**
**VPRSGKKRAAMMTEVLAHKIRAVEKKPIPRIKRKNNSTTTNNSKPSSLPTLGSNTETVQPEVKSETEPH**
**FILKSSDNTKTYSLMPSAPHPVKEASPGFSWSPKTASATPAPLKNDATASCGFSERSSTPHCTMPSGR**
**LSGANAAAADGPGISQLGEVAPLPTLSAPVMEPLINSEPSTGVTEPLTPHQPNHQPSFLTSPQDLASSP**
**MEEDEQHSEADEPPSDEPLSDDPLSPAEEKLPHIDEYWSDSEHIFLDANIGGVAIAPAHGSVLIECARRE**
**LHATTPVEHPNRNHPTRLSLVFYQHKNLNKPQHGFELNKIKFEAKEAKNKKMKASEQKDQAANEGPE**
**QSSEVNELNQIPSHKALTLTHDNVVTVSPYALTHVAGPYNHWV**ID

Name: **TALE_A-TET1(CD)**

**Keys:** NLS, TALE_A **TET1(1418-2136)**, His tag

MHHHHHHIDGGGGSDPKKKRKVDPKKKRKVDPKKKRKVGSTGSRNDGGGGSGGGGSGGGGSGRAV
DLRTLGYSQQQQEKIKPKVRSTVAQHHEALVGHGFTHAHIVALSQHPAALGTVAVTYQHIITALPEATHED
IVGVGKQWSGARALEALLTDAGELRGPPLQLDTGQLVKIAKRGGVTAMEAVHASRNALTGAPLNLTPDQ
VVAIAS**NN**GGKQALETVQRLLPVLCQDHGLTPDQVVAIAS**NI**GGKQALETVQRLLPVLCQDHGLTPDQVV
AIAS**NN**GGKQALETVQRLLPVLCQDHGLTPDQVVAIAS**NI**GGKQALETVQRLLPVLCQDHGLTPDQVVAIA
S**NI**GGKQALETVQRLLPVLCQDHGLTPDQVVAIAS**NI**GGKQALETVQRLLPVLCQDHGLTPDQVVAIAS**N**
**G**GGKQALETVQRLLPVLCQDHGLTPDQVVAIAS**NG**GGKQALETVQRLLPVLCQDHGLTPDQVVAIAS**NG**
GGKQALETVQRLLPVLCQDHGLTPDQVVAIAS**NN**GGKQALETVQRLLPVLCQDHGLTPDQVVAIAS**NI**GG
KQALETVQRLLPVLCQDHGLTPDQVVAIAS**HD**GGKQALETVQRLLPVLCQDHGLTPDQVVAIAS**NG**GGK
QALETVQRLLPVLCQDHGLTPDQVVAIAS**NN**GGKQALETVQRLLPVLCQDHGLTPDQVVAIAS**NN**GGKQ
ALETVQRLLPVLCQDHGLTPDQVVAIAS**HD**GGKQALETVQRLLPVLCQDHGLTPDQVVAIAS**NI**GGKQAL
ETVQRLLPVLCQDHGLTPDQVVAIAS**NG**GGKQALESIVAQLSRPDPALAALTNDHLVALACLGGRPAMDA
VKKGLPHAPELIRRVNRRIGERTSHRVARDPKKKRKVDPKKKRKVGGRGGGGSGGGGSGGGGSGPA**E**
**LPTCSCLDRVIQKDKGPYYTHLGAGPSVAAVREIMENRYGQKGNAIRIEIVVYTGKEGKSSHGCPIAKW**
**VLRRSSDEEKVLCLVRQRTGHHCPTAVMVVLIMVWDGIPLPMADRLYTELTENLKSYNGHPTDRRCTL**
**NENRTCTCQGIDPETCGASFSFGCSWSMYFNGCKFGRSPSPRRFRIDPSSPLHEKNLEDNLQSLATRL**
**APIYKQYAPVAYQNQVEYENVARECRLGSKEGRPFSGVTACLDFCAHPHRDIHNMNNGSTVVCTLTRE**
**DNRSLGVIPQDEQLHVLPLYKLSDTDEFGSKEGMEAKIKSGAIEVLAPRRKKRTCFTQPVPRSGKKRAA**
**MMTEVLAHKIRAVEKKPIPRIKRKNNSTTTNNSKPSSLPTLGSNTETVQPEVKSETEPHFILKSSDNTKTY**
**SLMPSAPHPVKEASPGFSWSPKTASATPAPLKNDATASCGFSERSSTPHCTMPSGRLSGANAAAADG**
**PGISQLGEVAPLPTLSAPVMEPLINSEPSTGVTEPLTPHQPNHQPSFLTSPQDLASSPMEEDEQHSEAD**
**EPPSDEPLSDDPLSPAEEKLPHIDEYWSDSEHIFLDANIGGVAIAPAHGSVLIECARRELHATTPVEHPN**
**RNHPTRLSLVFYQHKNLNKPQHGFELNKIKFEAKEAKNKKMKASEQKDQAANEGPEQSSEVNELNQIP**
**SHKALTLTHDNVVTVSPYALTHVAGPYNHWV**ID

Name: **TALE_B-TET1(CD)**

**Keys:** NLS, TALE-B **TET1(1418-2136),** His tag

MHHHHHHIDGGGGSDPKKKRKVDPKKKRKVDPKKKRKVGSTGSRNDGGGGSGGGGSGGGGSGRAV
DLRTLGYSQQQQEKIKPKVRSTVAQHHEALVGHGFTHAHIVALSQHPAALGTVAVTYQHIITALPEATHED
IVGVGKQWSGARALEALLTDAGELRGPPLQLDTGQLVKIAKRGGVTAMEAVHASRNALTGAPLNLTPDQ
VVAIAS**NI**GGKQALETVQRLLPVLCQDHGLTPDQVVAIAS**NN**GGKQALETVQRLLPVLCQDHGLTPDQVV
AIAS**HD**GGKQALETVQRLLPVLCQDHGLTPDQVVAIAS**NN**GGKQALETVQRLLPVLCQDHGLTPDQVVAI
AS**NN**GGKQALETVQRLLPVLCQDHGLTPDQVVAIAS**NN**GGKQALETVQRLLPVLCQDHGLTPDQVVAIA
S**HD**GGKQALETVQRLLPVLCQDHGLTPDQVVAIAS**NI**GGKQALETVQRLLPVLCQDHGLTPDQVVAIAS**N**
**N**GGKQALETVQRLLPVLCQDHGLTPDQVVAIAS**NG**GGKQALETVQRLLPVLCQDHGLTPDQVVAIAS**NI**G
GKQALETVQRLLPVLCQDHGLTPDQVVAIAS**NN**GGKQALETVQRLLPVLCQDHGLTPDQVVAIAS**HD**GG
KQALETVQRLLPVLCQDHGLTPDQVVAIAS**HD**GGKQALETVQRLLPVLCQDHGLTPDQVVAIAS**NN**GGK
QALETVQRLLPVLCQDHGLTPDQVVAIAS**HD**GGKQALETVQRLLPVLCQDHGLTPDQVVAIAS**NG**GGKQ
ALETVQRLLPVLCQDHGLTPDQVVAIAS**NG**GGKQALESIVAQLSRPDPALAALTNDHLVALACLGGRPAM
DAVKKGLPHAPELIRRVNRRIGERTSHRVARDPKKKRKVDPKKKRKVGGRGGGGSGGGGSGGGGSGP
A**ELPTCSCLDRVIQKDKGPYYTHLGAGPSVAAVREIMENRYGQKGNAIRIEIVVYTGKEGKSSHGCPIAK**
**WVLRRSSDEEKVLCLVRQRTGHHCPTAVMVVLIMVWDGIPLPMADRLYTELTENLKSYNGHPTDRRCT**
**LNENRTCTCQGIDPETCGASFSFGCSWSMYFNGCKFGRSPSPRRFRIDPSSPLHEKNLEDNLQSLATR**
**LAPIYKQYAPVAYQNQVEYENVARECRLGSKEGRPFSGVTACLDFCAHPHRDIHNMNNGSTVVCTLTR**
**EDNRSLGVIPQDEQLHVLPLYKLSDTDEFGSKEGMEAKIKSGAIEVLAPRRKKRTCFTQPVPRSGKKRA**
**AMMTEVLAHKIRAVEKKPIPRIKRKNNSTTTNNSKPSSLPTLGSNTETVQPEVKSETEPHFILKSSDNTK**
**TYSLMPSAPHPVKEASPGFSWSPKTASATPAPLKNDATASCGFSERSSTPHCTMPSGRLSGANAAAA**
**DGPGISQLGEVAPLPTLSAPVMEPLINSEPSTGVTEPLTPHQPNHQPSFLTSPQDLASSPMEEDEQHSE**
**ADEPPSDEPLSDDPLSPAEEKLPHIDEYWSDSEHIFLDANIGGVAIAPAHGSVLIECARRELHATTPVEH**
**PNRNHPTRLSLVFYQHKNLNKPQHGFELNKIKFEAKEAKNKKMKASEQKDQAANEGPEQSSEVNELN**
**QIPSHKALTLTHDNVVTVSPYALTHVAGPYNHWV**ID

Name: **TALE_D-TET1(CD)**

**Keys:** NLS, TALE_D **TET1(1418-2136),** His tag

MHHHHHHIDGGGGSDPKKKRKVDPKKKRKVDPKKKRKVGSTGSRNDGGGGSGGGGSGGGGSGRAV
DLRTLGYSQQQQEKIKPKVRSTVAQHHEALVGHGFTHAHIVALSQHPAALGTVAVTYQHIITALPEATHED
IVGVGKQWSGARALEALLTDAGELRGPPLQLDTGQLVKIAKRGGVTAMEAVHASRNALTGAPLNLTPDQ
VVAIAS**NI**GGKQALETVQRLLPVLCQDHGLTPDQVVAIAS**NI**GGKQALETVQRLLPVLCQDHGLTPDQVVA
IAS**NI**GGKQALETVQRLLPVLCQDHGLTPDQVVAIAS**NI**GGKQALETVQRLLPVLCQDHGLTPDQVVAIAS
**NI**GGKQALETVQRLLPVLCQDHGLTPDQVVAIAS**HD**GGKQALETVQRLLPVLCQDHGLTPDQVVAIAS**NN**
GGKQALETVQRLLPVLCQDHGLTPDQVVAIAS**NI**GGKQALETVQRLLPVLCQDHGLTPDQVVAIAS**NI**GG
KQALETVQRLLPVLCQDHGLTPDQVVAIAS**HD**GGKQALETVQRLLPVLCQDHGLTPDQVVAIAS**HD**GGK
QALETVQRLLPVLCQDHGLTPDQVVAIAS**NI**GGKQALETVQRLLPVLCQDHGLTPDQVVAIAS**NI**GGKQAL
ETVQRLLPVLCQDHGLTPDQVVAIAS**NG**GGKQALETVQRLLPVLCQDHGLTPDQVVAIAS**NI**GGKQALET
VQRLLPVLCQDHGLTPDQVVAIAS**NN**GGKQALETVQRLLPVLCQDHGLTPDQVVAIAS**NN**GGKQALETV
QRLLPVLCQDHGLTPDQVVAIAS**NI**GGKQALESIVAQLSRPDPALAALTNDHLVALACLGGRPAMDAVKK
GLPHAPELIRRVNRRIGERTSHRVARDPKKKRKVDPKKKRKVGGRGGGGSGGGGSGGGGSGPA**ELPT**
**CSCLDRVIQKDKGPYYTHLGAGPSVAAVREIMENRYGQKGNAIRIEIVVYTGKEGKSSHGCPIAKWVLR**
**RSSDEEKVLCLVRQRTGHHCPTAVMVVLIMVWDGIPLPMADRLYTELTENLKSYNGHPTDRRCTLNEN**
**RTCTCQGIDPETCGASFSFGCSWSMYFNGCKFGRSPSPRRFRIDPSSPLHEKNLEDNLQSLATRLAPIY**
**KQYAPVAYQNQVEYENVARECRLGSKEGRPFSGVTACLDFCAHPHRDIHNMNNGSTVVCTLTREDNR**
**SLGVIPQDEQLHVLPLYKLSDTDEFGSKEGMEAKIKSGAIEVLAPRRKKRTCFTQPVPRSGKKRAAMM**
**TEVLAHKIRAVEKKPIPRIKRKNNSTTTNNSKPSSLPTLGSNTETVQPEVKSETEPHFILKSSDNTKTYSL**
**MPSAPHPVKEASPGFSWSPKTASATPAPLKNDATASCGFSERSSTPHCTMPSGRLSGANAAAADGP**
**GISQLGEVAPLPTLSAPVMEPLINSEPSTGVTEPLTPHQPNHQPSFLTSPQDLASSPMEEDEQHSEADE**
**PPSDEPLSDDPLSPAEEKLPHIDEYWSDSEHIFLDANIGGVAIAPAHGSVLIECARRELHATTPVEHPNR**
**NHPTRLSLVFYQHKNLNKPQHGFELNKIKFEAKEAKNKKMKASEQKDQAANEGPEQSSEVNELNQIPS**
**HKALTLTHDNVVTVSPYALTHVAGPYNHWV**ID

Name: **TALE_F-TET1(CD)**

**Keys:** NLS, TALE_F **TET1(1418-2136)**, His tag

MHHHHHHIDGGGGSDPKKKRKVDPKKKRKVDPKKKRKVGSTGSRNDGGGGSGGGGSGGGGSGRAV
DLRTLGYSQQQQEKIKPKVRSTVAQHHEALVGHGFTHAHIVALSQHPAALGTVAVTYQHIITALPEATHED
IVGVGKQWSGARALEALLTDAGELRGPPLQLDTGQLVKIAKRGGVTAMEAVHASRNALTGAPLNLTPDQ
VVAIAS**NN**GGKQALETVQRLLPVLCQDHGLTPDQVVAIAS**NI**GGKQALETVQRLLPVLCQDHGLTPDQVV
AIAS**HD**GGKQALETVQRLLPVLCQDHGLTPDQVVAIAS**NN**GGKQALETVQRLLPVLCQDHGLTPDQVVAI
AS**HD**GGKQALETVQRLLPVLCQDHGLTPDQVVAIAS**NI**GGKQALETVQRLLPVLCQDHGLTPDQVVAIAS
**NN**GGKQALETVQRLLPVLCQDHGLTPDQVVAIAS**NI**GGKQALETVQRLLPVLCQDHGLTPDQVVAIAS**HD**
GGKQALETVQRLLPVLCQDHGLTPDQVVAIAS**NN**GGKQALETVQRLLPVLCQDHGLTPDQVVAIAS**HD**G
GKQALETVQRLLPVLCQDHGLTPDQVVAIAS**NG**GGKQALETVQRLLPVLCQDHGLTPDQVVAIAS**HD**GG
KQALETVQRLLPVLCQDHGLTPDQVVAIAS**HD**GGKQALETVQRLLPVLCQDHGLTPDQVVAIAS**NI**GGKQ
ALETVQRLLPVLCQDHGLTPDQVVAIAS**HD**GGKQALETVQRLLPVLCQDHGLTPDQVVAIAS**HD**GGKQAL
ETVQRLLPVLCQDHGLTPDQVVAIAS**NI**GGKQALESIVAQLSRPDPALAALTNDHLVALACLGGRPAMDA
VKKGLPHAPELIRRVNRRIGERTSHRVARDPKKKRKVDPKKKRKVGGRGGGGSGGGGSGGGGSGPA**E**
**LPTCSCLDRVIQKDKGPYYTHLGAGPSVAAVREIMENRYGQKGNAIRIEIVVYTGKEGKSSHGCPIAKW**
**VLRRSSDEEKVLCLVRQRTGHHCPTAVMVVLIMVWDGIPLPMADRLYTELTENLKSYNGHPTDRRCTL**
**NENRTCTCQGIDPETCGASFSFGCSWSMYFNGCKFGRSPSPRRFRIDPSSPLHEKNLEDNLQSLATRL**
**APIYKQYAPVAYQNQVEYENVARECRLGSKEGRPFSGVTACLDFCAHPHRDIHNMNNGSTVVCTLTRE**
**DNRSLGVIPQDEQLHVLPLYKLSDTDEFGSKEGMEAKIKSGAIEVLAPRRKKRTCFTQPVPRSGKKRAA**
**MMTEVLAHKIRAVEKKPIPRIKRKNNSTTTNNSKPSSLPTLGSNTETVQPEVKSETEPHFILKSSDNTKTY**
**SLMPSAPHPVKEASPGFSWSPKTASATPAPLKNDATASCGFSERSSTPHCTMPSGRLSGANAAAADG**
**PGISQLGEVAPLPTLSAPVMEPLINSEPSTGVTEPLTPHQPNHQPSFLTSPQDLASSPMEEDEQHSEAD**
**EPPSDEPLSDDPLSPAEEKLPHIDEYWSDSEHIFLDANIGGVAIAPAHGSVLIECARRELHATTPVEHPN**
**RNHPTRLSLVFYQHKNLNKPQHGFELNKIKFEAKEAKNKKMKASEQKDQAANEGPEQSSEVNELNQIP**
**SHKALTLTHDNVVTVSPYALTHVAGPYNHWV**ID

Name: **PUFa-dTET1(CD) (dead TET1(CD))**

**Keys:** NLS, PUFa, dTET1(1418-2136 (**H1671Y, D1673A**))

MIDGGGGSDPKKKRKVDPKKKRKVDPKKKRKVGSTGSRNDGGGGSGGGGSGGGGSGRAGILPPKKKR
KVSRGRSRLLEDFRNNRYPNLQLREIAGHIMEFSQDQHGSRFIQLKLERATPAERQLVFNEILQAAYQLM
VDVFGNYVIQKFFEFGSLEQKLALAERIRGHVLSLALQMYGSRVIEKALEFIPSDQQNEMVRELDGHVLKC
VKDQNGNHVVQKCIECVQPQSLQFIIDAFKGQVFALSTHPYGCRVIQRILEHCLPDQTLPILEELHQHTEQL
VQDQYGNYVIQHVLEHGRPEDKSKIVAEIRGNVLVLSQHKFASNVVEKCVTHASRTERAVLIDEVCTMND
GPHSALYTMMKDQYANYVVQKMIDVAEPGQRKIVMHKIRPHIATLRKYTYGKHILAKLEKYYMKNGVDLG
DPKKKRKVDPKKKRKVGGRGGGGSGGGGSGGGGSGPAELPTCSCLDRVIQKDKGPYYTHLGAGPSVA
AVREIMENRYGQKGNAIRIEIVVYTGKEGKSSHGCPIAKWVLRRSSDEEKVLCLVRQRTGHHCPTAVMVV
LIMVWDGIPLPMADRLYTELTENLKSYNGHPTDRRCTLNENRTCTCQGIDPETCGASFSFGCSWSMYFN
GCKFGRSPSPRRFRIDPSSPLHEKNLEDNLQSLATRLAPIYKQYAPVAYQNQVEYENVARECRLGSKEG
RPFSGVTACLDFCAHP**Y**R**A**IHNMNNGSTVVCTLTREDNRSLGVIPQDEQLHVLPLYKLSDTDEFGSKEG
MEAKIKSGAIEVLAPRRKKRTCFTQPVPRSGKKRAAMMTEVLAHKIRAVEKKPIPRIKRKNNSTTTNNSKP
SSLPTLGSNTETVQPEVKSETEPHFILKSSDNTKTYSLMPSAPHPVKEASPGFSWSPKTASATPAPLKND
ATASCGFSERSSTPHCTMPSGRLSGANAAAADGPGISQLGEVAPLPTLSAPVMEPLINSEPSTGVTEPLT
PHQPNHQPSFLTSPQDLASSPMEEDEQHSEADEPPSDEPLSDDPLSPAEEKLPHIDEYWSDSEHIFLDA
NIGGVAIAPAHGSVLIECARRELHATTPVEHPNRNHPTRLSLVFYQHKNLNKPQHGFELNKIKFEAKEAKN
KKMKASEQKDQAANEGPEQSSEVNELNQIPSHKALTLTHDNVVTVSPYALTHVAGPYNHWVID

Name: **PUFa-hGADD45A-TET1(CD)**

**Keys:** NLS, PUFa, GADD45A, **TET1(1418-2136)**

MIDGGGGSDPKKKRKVDPKKKRKVDPKKKRKVGSTGSRNDGGGGSGGGGSGGGGSGRAGILPPKKKR
KVSRGRSRLLEDFRNNRYPNLQLREIAGHIMEFSQDQHGSRFIQLKLERATPAERQLVFNEILQAAYQLM
VDVFGNYVIQKFFEFGSLEQKLALAERIRGHVLSLALQMYGSRVIEKALEFIPSDQQNEMVRELDGHVLKC

| |
|---|
| VKDQNGNHVVQKCIECVQPQSLQFIIDAFKGQVFALSTHPYGCRVIQRILEHCLPDQTLPILEELHQHTEQL<br>VQDQYGNYVIQHVLEHGRPEDKSKIVAEIRGNVLVLSQHKFASNVVEKCVTHASRTERAVLIDEVCTMND<br>GPHSALYTMMKDQYANYVVQKMIDVAEPGQRKIVMHKIRPHIATLRKYTYGKHILAKLEKYYMKNGVDLG<br>DPKKKRKVDPKKKRKVGGRGGGSGGGSGGGSGGGSGGGSGGGSLTLEEFSAGEQKTERMDKVGDAL<br>EEVLSKALSQRTITVGVYEAAKLLNVDPDNVVLCLLAADEDDDRDVALQIHFTLIQAFCCENDINILRVSNP<br>GRLAELLLLETDAGPAASEGAEQPPDLHCVLVTNPHSSQWKDPALSQLICFCRESRYMDQWVPVINLPE<br>RSRGRGGGGSGGGGSGGGGSGPA**ELPTCSCLDRVIQKDKGPYYTHLGAGPSVAAVREIMENRYGQK**<br>**GNAIRIEIVVYTGKEGKSSHGCPIAKWVLRRSSDEEKVLCLVRQRTGHHCPTAVMVVLIMVWDGIPLPM**<br>**ADRLYTELTENLKSYNGHPTDRRCTLNENRTCTCQGIDPETCGASFSFGCSWSMYFNGCKFGRSPSPR**<br>**RFRIDPSSPLHEKNLEDNLQSLATRLAPIYKQYAPVAYQNQVEYENVARECRLGSKEGRPFSGVTACLD**<br>**FCAHPHRDIHNMNNGSTVVCTLTREDNRSLGVIPQDEQLHVLPLYKLSDTDEFGSKEGMEAKIKSGAIE**<br>**VLAPRRKKRTCFTQPVPRSGKKRAAMMTEVLAHKIRAVEKKPIPRIKRKNNSTTTNNSKPSSLPTLGSN**<br>**TETVQPEVKSETEPHFILKSSDNTKTYSLMPSAPHPVKEASPGFSWSPKTASATPAPLKNDATASCGFS**<br>**ERSSTPHCTMPSGRLSGANAAAADGPGISQLGEVAPLPTLSAPVMEPLINSEPSTGVTEPLTPHQPNH**<br>**QPSFLTSPQDLASSPMEEDEQHSEADEPPSDEPLSDDPLSPAEEKLPHIDEYWSDSEHIFLDANIGGVAI**<br>**APAHGSVLIECARRELHATTPVEHPNRNHPTRLSLVFYQHKNLNKPQHGFELNKIKFEAKEAKNKKMK**<br>**ASEQKDQAANEGPEQSSEVNELNQIPSHKALTLTHDNVVTVSPYALTHVAGPYNHWV**ID |
| Name: **hGADD45A-PUFa-TET1(CD)** |
| **Keys:** NLS, PUFa, GADD45A, **TET1(1418-2136)** |
| MTLEEFSAGEQKTERMDKVGDALEEVLSKALSQRTITVGVYEAAKLLNVDPDNVVLCLLAADEDDDRDVA<br>LQIHFTLIQAFCCENDINILRVSNPGRLAELLLLETDAGPAASEGAEQPPDLHCVLVTNPHSSQWKDPALS<br>QLICFCRESRYMDQWVPVINLPERSRTGAATMIDGGGGSDPKKKRKVDPKKKRKVDPKKKRKVGSTGS<br>RNDGGGGSGGGGSGGGGSGRAGILPPKKKRKVSRGRSRLLEDFRNNRYPNLQLREIAGHIMEFSQDQH<br>GSRFIQLKLERATPAERQLVFNEILQAAYQLMVDVFGNYVIQKFFEFGSLEQKLALAERIRGHVLSLALQM<br>YGSRVIEKALEFIPSDQQNEMVRELDGHVLKCVKDQNGNHVVQKCIECVQPQSLQFIIDAFKGQVFALST<br>HPYGCRVIQRILEHCLPDQTLPILEELHQHTEQLVQDQYGNYVIQHVLEHGRPEDKSKIVAEIRGNVLVLS<br>QHKFASNVVEKCVTHASRTERAVLIDEVCTMNDGPHSALYTMMKDQYANYVVQKMIDVAEPGQRKIVMH<br>KIRPHIATLRKYTYGKHILAKLEKYYMKNGVDLGDPKKKRKVDPKKKRKVGGRGGGGSGGGGSGGGGS<br>GPA**ELPTCSCLDRVIQKDKGPYYTHLGAGPSVAAVREIMENRYGQKGNAIRIEIVVYTGKEGKSSHGCPI**<br>**AKWVLRRSSDEEKVLCLVRQRTGHHCPTAVMVVLIMVWDGIPLPMADRLYTELTENLKSYNGHPTDRR**<br>**CTLNENRTCTCQGIDPETCGASFSFGCSWSMYFNGCKFGRSPSPRRFRIDPSSPLHEKNLEDNLQSLA**<br>**TRLAPIYKQYAPVAYQNQVEYENVARECRLGSKEGRPFSGVTACLDFCAHPHRDIHNMNNGSTVVCTL**<br>**TREDNRSLGVIPQDEQLHVLPLYKLSDTDEFGSKEGMEAKIKSGAIEVLAPRRKKRTCFTQPVPRSGKK**<br>**RAAMMTEVLAHKIRAVEKKPIPRIKRKNNSTTTNNSKPSSLPTLGSNTETVQPEVKSETEPHFILKSSDN**<br>**TKTYSLMPSAPHPVKEASPGFSWSPKTASATPAPLKNDATASCGFSERSSTPHCTMPSGRLSGANAA**<br>**AADGPGISQLGEVAPLPTLSAPVMEPLINSEPSTGVTEPLTPHQPNHQPSFLTSPQDLASSPMEEDEQH**<br>**SEADEPPSDEPLSDDPLSPAEEKLPHIDEYWSDSEHIFLDANIGGVAIAPAHGSVLIECARRELHATTPV**<br>**EHPNRNHPTRLSLVFYQHKNLNKPQHGFELNKIKFEAKEAKNKKMKASEQKDQAANEGPEQSSEVNE**<br>**LNQIPSHKALTLTHDNVVTVSPYALTHVAGPYNHWV**ID |
| Name: **PUFc-hGADD45A** |
| **Keys:** NLS, PUFc, GADD45A, HA tag |
| MIDGGGGSDPKKKRKVDPKKKRKVDPKKKRKVGSTGSRNDGGGGSGGGGSGGGGSGRAGILPPKKKR<br>KVSRGRSRLLEDFRNNRYPNLQLREIAGHIMEFSQDQHGSRFIQLKLERATPAERQLVFNEILQAAYQLM<br>VDVFGNYVIQKFFEFGSLEQKLALAERIRGHVLSLALQMYGSRVIEKALEFIPSDQQNEMVRELDGHVLKC<br>VKDQNGNHVVQKCIECVQPQSLQFIIDAFKGQVFALSTHPYGCRVIQRILEHCLPDQTLPILEELHQHTEQL<br>VQDQYGSYVIEHVLEHGRPEDKSKIVAEIRGNVLVLSQHKFANNVVQKCVTHASRTERAVLIDEVCTMND<br>GPHSALYTMMKDQYANYVVQKMIDVAEPGQRKIVMHKIRPHIATLRKYTYGKHILAKLEKYYMKNGVDLG<br>DPKKKRKVDPKKKRKVGGRGGGGSGGGGSGGGGSGPALTLEEFSAGEQKTERMDKVGDALEEVLSKA<br>LSQRTITVGVYEAAKLLNVDPDNVVLCLLAADEDDDRDVALQIHFTLIQAFCCENDINILRVSNPGRLAELLL<br>LETDAGPAASEGAEQPPDLHCVLVTNPHSSQWKDPALSQLICFCRESRYMDQWVPVINLPERSRYPYDV<br>PDYA |

Name: **hGADD45A-PUFc**

**Keys:** NLS, PUFc, GADD45A, HA tag

MTLEEFSAGEQKTERMDKVGDALEEVLSKALSQRTITVGVYEAAKLLNVDPDNVVLCLLAADEDDDRDVALQIHFTLIQAFCCENDINILRVSNPGRLAELLLLETDAGPAASEGAEQPPDLHCVLVTNPHSSQWKDPALSQLICFCRESRYMDQWVPVINLPERSRYPYDVPDYAIDGGGGSDPKKKRKVDPKKKRKVDPKKKRKVGSTGSRNDGGGGSGGGGSGGGGSGRAGILPPKKKRKVSRGRSRLLEDFRNNRYPNLQLREIAGHIMEFSQDQHGSRFIQLKLERATPAERQLVFNEILQAAYQLMVDVFGNYVIQKFFEFGSLEQKLALAERIRGHVLSLALQMYGSRVIEKALEFIPSDQQNEMVRELDGHVLKCVKDQNGNHVVQKCIECVQPQSLQFIIDAFKGQVFALSTHPYGCRVIQRILEHCLPDQTLPILEELHQHTEQLVQDQYGSYVIEHVLEHGRPEDKSKIVAEIRGNVLVLSQHKFANNVVQKCVTHASRTERAVLIDEVCTMNDGPHSALYTMMKDQYANYVVQKMIDVAEPGQRKIVMHKIRPHIATLRKYTYGKHILAKLEKYYMKNGVDLGDPKKKRKVDPKKKRKVGGRGGGGSGGGGSGGGGSGPA

Name: **hNEIL2-PUFa-TET1(CD)**

**Keys:** NLS, PUFa, NEIL2, **TET1(1418-2136)**, **Flag tag**

M**DYKDDDDK**PKKKRKLPEGPLVRKFHHLVSPFVGQQVVKTGGSSKKLQPASLQSLWLQDTQVHGKKLFLRFDLDEEMGPPGSSPTPEPPQKEVQKEGAADPKQVGEPSGQKTLDGSSRSAELVPQGEDDSEYLERDAPAGDAGRWLRVSFGLFGSVWVNDFSRAKKANKRGDWRDPSPRLVLHFGGGGFLAFYNCQLSWSSSPVVTPTCDILSEKFHRGQALEALGQAQPVCYTLLDQRYFSGLGNIIKNEALYRAGIHPLSLGSVLSASRREVLVDHVVEFSTAWLQGKFQGRPQHTQVYQKEQCPAGHQVMKEAFGPEDGLQRLTWWCPQCQPQLSEEPEQCQFSGAATMIDGGGGSDPKKKRKVDPKKKRKVDPKKKRKVGSTGSRNDGGGGSGGGGSGGGGSGRAGILPPKKKRKVSRGRSRLLEDFRNNRYPNLQLREIAGHIMEFSQDQHGSRFIQLKLERATPAERQLVFNEILQAAYQLMVDVFGNYVIQKFFEFGSLEQKLALAERIRGHVLSLALQMYGSRVIEKALEFIPSDQQNEMVRELDGHVLKCVKDQNGNHVVQKCIECVQPQSLQFIIDAFKGQVFALSTHPYGCRVIQRILEHCLPDQTLPILEELHQHTEQLVQDQYGNYVIQHVLEHGRPEDKSKIVAEIRGNVLVLSQHKFASNVVEKCVTHASRTERAVLIDEVCTMNDGPHSALYTMMKDQYANYVVQKMIDVAEPGQRKIVMHKIRPHIATLRKYTYGKHILAKLEKYYMKNGVDLGDPKKKRKVDPKKKRKVGGRGGGGSGGGGSGGGGSGPA**ELPTCSCLDRVIQKDKGPYYTHLGAGPSVAAVREIMENRYGQKGNAIRIEIVVYTGKEGKSSHGCPIAKWVLRRSSDEEKVLCLVRQRTGHHCPTAVMVVLIMVWDGIPLPMADRLYTELTENLKSYNGHPTDRRCTLNENRTCTCQGIDPETCGASFSFGCSWSMYFNGCKFGRSPSPRRFRIDPSSPLHEKNLEDNLQSLATRLAPIYKQYAPVAYQNQVEYENVARECRLGSKEGRPFSGVTACLDFCAHPHRDIHNMNNGSTVVCTLTREDNRSLGVIPQDEQLHVLPLYKLSDTDEFGSKEGMEAKIKSGAIEVLAPRRKKRTCFTQPVPRSGKKRAAMMTEVLAHKIRAVEKKPIPRIKRKNNSTTTNNSKPSSLPTLGSNTETVQPEVKSETEPHFILKSSDNTKTYSLMPSAPHPVKEASPGFSWSPKTASATPAPLKNDATASCGFSERSSTPHCTMPSGRLSGANAAAADGPGISQLGEVAPLPTLSAPVMEPLINSEPSTGVTEPLTPHQPNHQPSFLTSPQDLASSPMEEDEQHSEADEPPSDEPLSDDPLSPAEEKLPHIDEYWSDSEHIFLDANIGGVAIAPAHGSVLIECARRELHATTPVEHPNRNHPTRLSLVFYQHKNLNKPQHGFELNKIKFEAKEAKNKKMKASEQKDQAANEGPEQSSEVNELNQIPSHKALTLTHDNVVTVSPYALTHVAGPYNHWV**ID

Name: **PUFa-hNEIL2-TET1(CD)**

**Keys:** NLS, PUFa, NEIL2, **TET1(1418-2136)**

MIDGGGGSDPKKKRKVDPKKKRKVDPKKKRKVGSTGSRNDGGGGSGGGGSGGGGSGRAGILPPKKKRKVSRGRSRLLEDFRNNRYPNLQLREIAGHIMEFSQDQHGSRFIQLKLERATPAERQLVFNEILQAAYQLMVDVFGNYVIQKFFEFGSLEQKLALAERIRGHVLSLALQMYGSRVIEKALEFIPSDQQNEMVRELDGHVLKCVKDQNGNHVVQKCIECVQPQSLQFIIDAFKGQVFALSTHPYGCRVIQRILEHCLPDQTLPILEELHQHTEQLVQDQYGNYVIQHVLEHGRPEDKSKIVAEIRGNVLVLSQHKFASNVVEKCVTHASRTERAVLIDEVCTMNDGPHSALYTMMKDQYANYVVQKMIDVAEPGQRKIVMHKIRPHIATLRKYTYGKHILAKLEKYYMKNGVDLGDPKKKRKVDPKKKRKVGGRGGGSGGGSGGGSGGGSGGGSGGGSLPEGPLVRKFHHLVSPFVGQQVVKTGGSSKKLQPASLQSLWLQDTQVHGKKLFLRFDLDEEMGPPGSSPTPEPPQKEVQKEGAADPKQVGEPSGQKTLDGSSRSAELVPQGEDDSEYLERDAPAGDAGRWLRVSFGLFGSVWVNDFSRAKKANKRGDWRDPSPRLVLHFGGGGFLAFYNCQLSWSSSPVVTPTCDILSEKFHRGQALEALGQAQPVCYTLLDQRYFSGLGNIIKNEALYRAGIHPLSLGSVLSASRREVLVDHVVEFSTAWLQGKFQGRPQHTQVYQKEQCPAGHQ

| |
|---|
| VMKEAFGPEDGLQRLTWWCPQCQPQLSEEPEQCQFSRGGGGSGGGGSGGGGSGPA**ELPTCSCLDR<br>VIQKDKGPYYTHLGAGPSVAAVREIMENRYGQKGNAIRIEIVVYTGKEGKSSHGCPIAKWVLRRSSDEE<br>KVLCLVRQRTGHHCPTAVMVVLIMVWDGIPLPMADRLYTELTENLKSYNGHPTDRRCTLNENRTCTCQ<br>GIDPETCGASFSFGCSWSMYFNGCKFGRSPSPRRFRIDPSSPLHEKNLEDNLQSLATRLAPIYKQYAPV<br>AYQNQVEYENVARECRLGSKEGRPFSGVTACLDFCAHPHRDIHNMNNGSTVVCTLTREDNRSLGVIPQ<br>DEQLHVLPLYKLSDTDEFGSKEGMEAKIKSGAIEVLAPRRKKRTCFTQPVPRSGKKRAAMMTEVLAHKI<br>RAVEKKPIPRIKRKNNSTTTNNSKPSSLPTLGSNTETVQPEVKSETEPHFILKSSDNTKTYSLMPSAPHP<br>VKEASPGFSWSPKTASATPAPLKNDATASCGFSERSSTPHCTMPSGRLSGANAAAADGPGISQLGEV<br>APLPTLSAPVMEPLINSEPSTGVTEPLTPHQPNHQPSFLTSPQDLASSPMEEDEQHSEADEPPSDEPLS<br>DDPLSPAEEKLPHIDEYWSDSEHIFLDANIGGVAIAPAHGSVLIECARRELHATTPVEHPNRNHPTRLSL<br>VFYQHKNLNKPQHGFELNKIKFEAKEAKNKKMKASEQKDQAANEGPEQSSEVNELNQIPSHKALTLTH<br>DNVVTVSPYALTHVAGPYNHWV**ID |
| Name: **PUFc-hNEIL2** |
| **Keys:** NLS, PUFc, NEIL2 |
| MIDGGGGSDPKKKRKVDPKKKRKVDPKKKRKVGSTGSRNDGGGGSGGGGSGGGGSGRAGILPPKKKR<br>KVSRGRSRLLEDFRNNRYPNLQLREIAGHIMEFSQDQHGSRFIQLKLERATPAERQLVFNEILQAAYQLM<br>VDVFGNYVIQKFFEFGSLEQKLALAERIRGHVLSLALQMYGSRVIEKALEFIPSDQQNEMVRELDGHVLKC<br>VKDQNGNHVVQKCIECVQPQSLQFIIDAFKGQVFALSTHPYGCRVIQRILEHCLPDQTLPILEELHQHTEQL<br>VQDQYGSYVIEHVLEHGRPEDKSKIVAEIRGNVLVLSQHKFANNVVQKCVTHASRTERAVLIDEVCTMND<br>GPHSALYTMMKDQYANYVVQKMIDVAEPGQRKIVMHKIRPHIATLRKYTYGKHILAKLEKYYMKNGVDLG<br>DPKKKRKVDPKKKRKVGGRGGGGSGGGGSGGGGSGPALPEGPLVRKFHHLVSPFVGQQVVKTGGSS<br>KKLQPASLQSLWLQDTQVHGKKLFLRFDLDEEMGPPGSSPTPEPPQKEVQKEGAADPKQVGEPSGQKT<br>LDGSSRSAELVPQGEDDSEYLERDAPAGDAGRWLRVSFGLFGSVWVNDFSRAKKANKRGDWRDPSPR<br>LVLHFGGGGFLAFYNCQLSWSSSPVVTPTCDILSEKFHRGQALEALGQAQPVCYTLLDQRYFSGLGNIIK<br>NEALYRAGIHPLSLGSVLSASRREVLVDHVVEFSTAWLQGKFQGRPQHTQVYQKEQCPAGHQVMKEAF<br>GPEDGLQRLTWWCPQCQPQLSEEPEQCQFS |
| Name: **hNEIL2- PUFc** |
| **Keys:** NLS, PUFc, NEIL2 |
| MPEGPLVRKFHHLVSPFVGQQVVKTGGSSKKLQPASLQSLWLQDTQVHGKKLFLRFDLDEEMGPPGSS<br>PTPEPPQKEVQKEGAADPKQVGEPSGQKTLDGSSRSAELVPQGEDDSEYLERDAPAGDAGRWLRVSF<br>GLFGSVWVNDFSRAKKANKRGDWRDPSPRLVLHFGGGGFLAFYNCQLSWSSSPVVTPTCDILSEKFHR<br>GQALEALGQAQPVCYTLLDQRYFSGLGNIIKNEALYRAGIHPLSLGSVLSASRREVLVDHVVEFSTAWLQ<br>GKFQGRPQHTQVYQKEQCPAGHQVMKEAFGPEDGLQRLTWWCPQCQPQLSEEPEQCQFSIDGGGGS<br>DPKKKRKVDPKKKRKVDPKKKRKVGSTGSRNDGGGGSGGGGSGGGGSGRAGILPPKKKRKVSRGRSR<br>LLEDFRNNRYPNLQLREIAGHIMEFSQDQHGSRFIQLKLERATPAERQLVFNEILQAAYQLMVDVFGNYVI<br>QKFFEFGSLEQKLALAERIRGHVLSLALQMYGSRVIEKALEFIPSDQQNEMVRELDGHVLKCVKDQNGNH<br>VVQKCIECVQPQSLQFIIDAFKGQVFALSTHPYGCRVIQRILEHCLPDQTLPILEELHQHTEQLVQDQYGSY<br>VIEHVLEHGRPEDKSKIVAEIRGNVLVLSQHKFANNVVQKCVTHASRTERAVLIDEVCTMNDGPHSALYT<br>MMKDQYANYVVQKMIDVAEPGQRKIVMHKIRPHIATLRKYTYGKHILAKLEKYYMKNGVDLGDPKKKRKV<br>DPKKKRKVGGRGGGGSGGGGSGGGGSGPA |
| Name: **hNEIL1-PUFa-TET1(CD)** |
| **Keys:** NLS, PUFa, NEIL1, **TET1(1418-2136), Flag tag** |
| M**DYKDDDDK**PKKKRKLPEGPELHLASQFVNEACRALVFGGCVEKSSVSRNPEVPFESSAYRISASARGK<br>ELRLILSPLPGAQPQQEPLALVFRFGMSGSFQLVPREELPRHAHLRFYTAPPGPRLALCFVDIRRFGRWD<br>LGGKWQPGRGPCVLQEYQQFRENVLRNLADKAFDRPICEALLDQRFFNGIGNYLRAEILYRLKIPPFEKA<br>RSVLEALQQHRPSPELTLSQKIRTKLQNPDLLELCHSVPKEVVQLGGRGYGSESGEEDFAAFRAWLRCY<br>GMPGMSSLQDRHGRTIWFQGDPGPLAPKGRKSRKKKSKATQLSPEDRVEDALPPSKAPSRTRRAKRDL<br>PKRTATQRPEGTSLQQDPEAPTVPKKGRRKGRQAASGHCRPRKVKADIPSLEPEGTSASGAATMIDGG<br>GGSDPKKKRKVDPKKKRKVDPKKKRKVGSTGSRNDGGGGSGGGGSGGGGSGRAGILPPKKKRKVSR<br>GRSRLLEDFRNNRYPNLQLREIAGHIMEFSQDQHGSRFIQLKLERATPAERQLVFNEILQAAYQLMVDVF |

GNYVIQKFFEFGSLEQKLALAERIRGHVLSLALQMYGSRVIEKALEFIPSDQQNEMVRELDGHVLKCVKDQ
NGNHVVQKCIECVQPQSLQFIIDAFKGQVFALSTHPYGCRVIQRILEHCLPDQTLPILEELHQHTEQLVQD
QYGNYVIQHVLEHGRPEDKSKIVAEIRGNVLVLSQHKFASNVVEKCVTHASRTERAVLIDEVCTMNDGPH
SALYTMMKDQYANYVVQKMIDVAEPGQRKIVMHKIRPHIATLRKYTYGKHILAKLEKYYMKNGVDLGDPK
KKRKVDPKKKRKVGGRGGGGSGGGGSGGGGSGPA**ELPTCSCLDRVIQKDKGPYYTHLGAGPSVAAV
REIMENRYGQKGNAIRIEIVVYTGKEGKSSHGCPIAKWVLRRSSDEEKVLCLVRQRTGHHCPTAVMVVLI
MVWDGIPLPMADRLYTELTENLKSYNGHPTDRRCTLNENRTCTCQGIDPETCGASFSFGCSWSMYFNG
CKFGRSPSPRRFRIDPSSPLHEKNLEDNLQSLATRLAPIYKQYAPVAYQNQVEYENVARECRLGSKEG
RPFSGVTACLDFCAHPHRDIHNMNNGSTVVCTLTREDNRSLGVIPQDEQLHVLPLYKLSDTDEFGSKEG
MEAKIKSGAIEVLAPRRKKRTCFTQPVPRSGKKRAAMMTEVLAHKIRAVEKKPIPRIKRKNNSTTTNNS
KPSSLPTLGSNTETVQPEVKSETEPHFILKSSDNTKTYSLMPSAPHPVKEASPGFSWSPKTASATPAPL
KNDATASCGFSERSSTPHCTMPSGRLSGANAAAADGPGISQLGEVAPLPTLSAPVMEPLINSEPSTGV
TEPLTPHQPNHQPSFLTSPQDLASSPMEEDEQHSEADEPPSDEPLSDDPLSPAEEKLPHIDEYWSDSE
HIFLDANIGGVAIAPAHGSVLIECARRELHATTPVEHPNRNHPTRLSLVFYQHKNLNKPQHGFELNKIKF
EAKEAKNKKMKASEQKDQAANEGPEQSSEVNELNQIPSHKALTLTHDNVVTVSPYALTHVAGPYNHW
V**ID

Name: **PUFa-hNEIL1-TET1(CD)**

**Keys:** NLS, PUFa, NEIL1, **TET1(1418-2136), Flag tag**

MIDGGGGSDPKKKRKVDPKKKRKVDPKKKRKVGSTGSRNDGGGGSGGGGSGGGGSGRAGILPPKKKR
KVSRGRSRLLEDFRNNRYPNLQLREIAGHIMEFSQDQHGSRFIQLKLERATPAERQLVFNEILQAAYQLM
VDVFGNYVIQKFFEFGSLEQKLALAERIRGHVLSLALQMYGSRVIEKALEFIPSDQQNEMVRELDGHVLKC
VKDQNGNHVVQKCIECVQPQSLQFIIDAFKGQVFALSTHPYGCRVIQRILEHCLPDQTLPILEELHQHTEQL
VQDQYGNYVIQHVLEHGRPEDKSKIVAEIRGNVLVLSQHKFASNVVEKCVTHASRTERAVLIDEVCTMND
GPHSALYTMMKDQYANYVVQKMIDVAEPGQRKIVMHKIRPHIATLRKYTYGKHILAKLEKYYMKNGVDLG
DPKKKRKVDPKKKRKVGGRGGGSGGGSGGGSGGGSGGGSGGGSLPEGPELHLASQFVNEACRALVF
GGCVEKSSVSRNPEVPFESSAYRISASARGKELRLILSPLPGAQPQQEPLALVFRFGMSGSFQLVPREEL
PRHAHLRFYTAPPGPRLALCFVDIRRFGRWDLGGKWQPGRGPCVLQEYQQFRENVLRNLADKAFDRPI
CEALLDQRFFNGIGNYLRAEILYRLKIPPFEKARSVLEALQQHRPSPELTLSQKIRTKLQNPDLLELCHSVP
KEVVQLGGRGYGSESGEEDFAAFRAWLRCYGMPGMSSLQDRHGRTIWFQGDPGPLAPKGRKSRKKKS
KATQLSPEDRVEDALPPSKAPSRTRRAKRDLPKRTATQRPEGTSLQQDPEAPTVPKKGRRKGRQAASG
HCRPRKVKADIPSLEPEGTSASRGGGGSGGGGSGGGGSGPA**ELPTCSCLDRVIQKDKGPYYTHLGAG
PSVAAVREIMENRYGQKGNAIRIEIVVYTGKEGKSSHGCPIAKWVLRRSSDEEKVLCLVRQRTGHHCPT
AVMVVLIMVWDGIPLPMADRLYTELTENLKSYNGHPTDRRCTLNENRTCTCQGIDPETCGASFSFGCS
WSMYFNGCKFGRSPSPRRFRIDPSSPLHEKNLEDNLQSLATRLAPIYKQYAPVAYQNQVEYENVAREC
RLGSKEGRPFSGVTACLDFCAHPHRDIHNMNNGSTVVCTLTREDNRSLGVIPQDEQLHVLPLYKLSDTD
EFGSKEGMEAKIKSGAIEVLAPRRKKRTCFTQPVPRSGKKRAAMMTEVLAHKIRAVEKKPIPRIKRKNN
STTTNNSKPSSLPTLGSNTETVQPEVKSETEPHFILKSSDNTKTYSLMPSAPHPVKEASPGFSWSPKTA
SATPAPLKNDATASCGFSERSSTPHCTMPSGRLSGANAAAADGPGISQLGEVAPLPTLSAPVMEPLIN
SEPSTGVTEPLTPHQPNHQPSFLTSPQDLASSPMEEDEQHSEADEPPSDEPLSDDPLSPAEEKLPHIDE
YWSDSEHIFLDANIGGVAIAPAHGSVLIECARRELHATTPVEHPNRNHPTRLSLVFYQHKNLNKPQHGF
ELNKIKFEAKEAKNKKMKASEQKDQAANEGPEQSSEVNELNQIPSHKALTLTHDNVVTVSPYALTHVA
GPYNHWV**ID

Name: **hNEIL3-PUFa-TET1(CD)**

**Keys:** NLS, PUFa, NEIL3, **TET1(1418-2136), Flag tag**

M**DYKDDDDK**PKKKRKLVEGPGCTLNGEKIRARVLPGQAVTGVRGSALRSLQGRALRLAASTVVVSPQA
AALNNDSSQNVLSLFNGYVYSGVETLGKELFMYFGPKALRIHFGMKGFIMINPLEYKYKNGASPVLEVQLT
KDLICFFDSSVELRNSMESQQRIRMMKELDVCSPEFSFLRAESEVKKQKGRMLGDVLMDQNVLPGVGNII
KNEALFDSGLHPAVKVCQLTDEQIHHLMKMIRDFSILFYRCRKAGLALSKHYKVYKRPNCGQCHCRITVC
RFGDNNRMTYFCPHCQKENPQHVDICKLPTRNTIISWTSSRVDHVMDSVARKSEEHWTCVVCTLINKPS
SKACDACLTSRPIDSVLKSEENSTVFSHLMKYPCNTFGKPHTEVKINRKTAFGTTTLVLTDFSNKSSTLER
KTKQNQILDEEFQNSPPASVCLNDIQHPSKKTTNDITQLSSKVNISPTISSESKLFSPAHKKPKTAHYSSPE
LKSCNPGYSNSELQINMTDGPRTLNPDSPRCSKHNRLCILRVVRKDGENKGRQFYACPLPREAQCGFFE

WADLSFPFCNHGKRSTMKTVLKIGPNNGKNFFVCPLGKEKQCNFFQWAENGPGIKIIPGCGAATMIDGG
GGSDPKKKRKVDPKKKRKVDPKKKRKVGSTGSRNDGGGGSGGGGSGGGGSGRAGILPPKKKRKVSR
GRSRLLEDFRNNRYPNLQLREIAGHIMEFSQDQHGSRFIQLKLERATPAERQLVFNEILQAAYQLMVDVF
GNYVIQKFFEFGSLEQKLALAERIRGHVLSLALQMYGSRVIEKALEFIPSDQQNEMVRELDGHVLKCVKDQ
NGNHVVQKCIECVQPQSLQFIIDAFKGQVFALSTHPYGCRVIQRILEHCLPDQTLPILEELHQHTEQLVQD
QYGNYVIQHVLEHGRPEDKSKIVAEIRGNVLVLSQHKFASNVVEKCVTHASRTERAVLIDEVCTMNDGPH
SALYTMMKDQYANYVVQKMIDVAEPGQRKIVMHKIRPHIATLRKYTYGKHILAKLEKYYMKNGVDLGDPK
KKRKVDPKKKRKVGGRGGGGSGGGGSGGGGSGPA**ELPTCSCLDRVIQKDKGPYYTHLGAGPSVAAV**
**REIMENRYGQKGNAIRIEIVVYTGKEGKSSHGCPIAKWVLRRSSDEEKVLCLVRQRTGHHCPTAVMVVLI**
**MVWDGIPLPMADRLYTELTENLKSYNGHPTDRRCTLNENRTCTCQGIDPETCGASFSFGCSWSMYFNG**
**CKFGRSPSPRRFRIDPSSPLHEKNLEDNLQSLATRLAPIYKQYAPVAYQNQVEYENVARECRLGSKEG**
**RPFSGVTACLDFCAHPHRDIHNMNNGSTVVCTLTREDNRSLGVIPQDEQLHVLPLYKLSDTDEFGSKEG**
**MEAKIKSGAIEVLAPRRKKRTCFTQPVPRSGKKRAAMMTEVLAHKIRAVEKKPIPRIKRKNNSTTTNNS**
**KPSSLPTLGSNTETVQPEVKSETEPHFILKSSDNTKTYSLMPSAPHPVKEASPGFSWSPKTASATPAPL**
**KNDATASCGFSERSSTPHCTMPSGRLSGANAAAADGPGISQLGEVAPLPTLSAPVMEPLINSEPSTGV**
**TEPLTPHQPNHQPSFLTSPQDLASSPMEEDEQHSEADEPPSDEPLSDDPLSPAEEKLPHIDEYWSDSE**
**HIFLDANIGGVAIAPAHGSVLIECARRELHATTPVEHPNRNHPTRLSLVFYQHKNLNKPQHGFELNKIKF**
**EAKEAKNKKMKASEQKDQAANEGPEQSSEVNELNQIPSHKALTLTHDNVVTVSPYALTHVAGPYNHW**
**V**ID

Name: **hTDG-PUFa-TET1(CD)**

**Keys:** NLS, PUFa, TDG, **TET1(1418-2136), Flag tag**

M**DYKDDDDK**PKKKRKLEAENAGSYSLQQAQAFYTFPFQQLMAEAPNMAVVNEQQMPEEVPAPAPAQE
PVQEAPKGRKRKPRTTEPKQPVEPKKPVESKKSGKSAKSKEKQEKITDTFKVKRKVDRFNGVSEAELLTK
TLPDILTFNLDIVIIGINPGLMAAYKGHHYPGPGNHFWKCLFMSGLSEVQLNHMDDHTLPGKYGIGFTNMV
ERTTPGSKDLSSKEFREGGRILVQKLQKYQPRIAVFNGKCIYEIFSKEVFGVKVKNLEFGLQPHKIPDTETL
CYVMPSSSARCAQFPRAQDKVHYYIKLKDLRDQLKGIERNMDVQEVQYTFDLQLAQEDAKKMAVKEEKY
DPGYEAAYGGAYGENPCSSEPCGFSSNGLIESVELRGESAFSGIPNGQWMTQSFTDQIPSFSNHCGTQ
EQEEESHATGAATMIDGGGGSDPKKKRKVDPKKKRKVDPKKKRKVGSTGSRNDGGGGSGGGGSGGG
GSGRAGILPPKKKRKVSRGRSRLLEDFRNNRYPNLQLREIAGHIMEFSQDQHGSRFIQLKLERATPAERQ
LVFNEILQAAYQLMVDVFGNYVIQKFFEFGSLEQKLALAERIRGHVLSLALQMYGSRVIEKALEFIPSDQQN
EMVRELDGHVLKCVKDQNGNHVVQKCIECVQPQSLQFIIDAFKGQVFALSTHPYGCRVIQRILEHCLPDQ
TLPILEELHQHTEQLVQDQYGNYVIQHVLEHGRPEDKSKIVAEIRGNVLVLSQHKFASNVVEKCVTHASRT
ERAVLIDEVCTMNDGPHSALYTMMKDQYANYVVQKMIDVAEPGQRKIVMHKIRPHIATLRKYTYGKHILAK
LEKYYMKNGVDLGDPKKKRKVDPKKKRKVGGRGGGGSGGGGSGGGGSGPA**ELPTCSCLDRVIQKDK**
**GPYYTHLGAGPSVAAVREIMENRYGQKGNAIRIEIVVYTGKEGKSSHGCPIAKWVLRRSSDEEKVLCLV**
**RQRTGHHCPTAVMVVLIMVWDGIPLPMADRLYTELTENLKSYNGHPTDRRCTLNENRTCTCQGIDPET**
**CGASFSFGCSWSMYFNGCKFGRSPSPRRFRIDPSSPLHEKNLEDNLQSLATRLAPIYKQYAPVAYQNQ**
**VEYENVARECRLGSKEGRPFSGVTACLDFCAHPHRDIHNMNNGSTVVCTLTREDNRSLGVIPQDEQLH**
**VLPLYKLSDTDEFGSKEGMEAKIKSGAIEVLAPRRKKRTCFTQPVPRSGKKRAAMMTEVLAHKIRAVEK**
**KPIPRIKRKNNSTTTNNSKPSSLPTLGSNTETVQPEVKSETEPHFILKSSDNTKTYSLMPSAPHPVKEAS**
**PGFSWSPKTASATPAPLKNDATASCGFSERSSTPHCTMPSGRLSGANAAAADGPGISQLGEVAPLPTL**
**SAPVMEPLINSEPSTGVTEPLTPHQPNHQPSFLTSPQDLASSPMEEDEQHSEADEPPSDEPLSDDPLSP**
**AEEKLPHIDEYWSDSEHIFLDANIGGVAIAPAHGSVLIECARRELHATTPVEHPNRNHPTRLSLVFYQHK**
**NLNKPQHGFELNKIKFEAKEAKNKKMKASEQKDQAANEGPEQSSEVNELNQIPSHKALTLTHDNVVTV**
**SPYALTHVAGPYNHWV**ID

Name: **PUFa-TDG-TET1(CD)**

**Keys:** NLS, PUFa, TDG **TET1(1418-2136), Flag tag**

MIDGGGGSDPKKKRKVDPKKKRKVDPKKKRKVGSTGSRNDGGGGSGGGGSGGGGSGRAGILPPKKKR
KVSRGRSRLLEDFRNNRYPNLQLREIAGHIMEFSQDQHGSRFIQLKLERATPAERQLVFNEILQAAYQLM
VDVFGNYVIQKFFEFGSLEQKLALAERIRGHVLSLALQMYGSRVIEKALEFIPSDQQNEMVRELDGHVLKC
VKDQNGNHVVQKCIECVQPQSLQFIIDAFKGQVFALSTHPYGCRVIQRILEHCLPDQTLPILEELHQHTEQL
VQDQYGNYVIQHVLEHGRPEDKSKIVAEIRGNVLVLSQHKFASNVVEKCVTHASRTERAVLIDEVCTMND

GPHSALYTMMKDQYANYVVQKMIDVAEPGQRKIVMHKIRPHIATLRKYTYGKHILAKLEKYYMKNGVDLG
DPKKKRKVDPKKKRKVGGRGGGSGGGSGGGSGGGSGGGSGGGSLEAENAGSYSLQQAQAFYTFPFQ
QLMAEAPNMAVVNEQQMPEEVPAPAPAQEPVQEAPKGRKRKPRTTEPKQPVEPKKPVESKKSGKSAKS
KEKQEKITDTFKVKRKVDRFNGVSEAELLTKTLPDILTFNLDIVIIGINPGLMAAYKGHHYPGPGNHFWKCL
FMSGLSEVQLNHMDDHTLPGKYGIGFTNMVERTTPGSKDLSSKEFREGGRILVQKLQKYQPRIAVFNGK
CIYEIFSKEVFGVKVKNLEFGLQPHKIPDTETLCYVMPSSSARCAQFPRAQDKVHYYIKLKDLRDQLKGIER
NMDVQEVQYTFDLQLAQEDAKKMAVKEEKYDPGYEAAYGGAYGENPCSSEPCGFSSNGLIESVELRGE
SAFSGIPNGQWMTQSFTDQIPSFSNHCGTQEQEEESHAGRGGGGSGGGGSGGGGSGPA**ELPTCSCL**
**DRVIQKDKGPYYTHLGAGPSVAAVREIMENRYGQKGNAIRIEIVVYTGKEGKSSHGCPIAKWVLRRSSD**
**EEKVLCLVRQRTGHHCPTAVMVVLIMVWDGIPLPMADRLYTELTENLKSYNGHPTDRRCTLNENRTCT**
**CQGIDPETCGASFSFGCSWSMYFNGCKFGRSPSPRRFRIDPSSPLHEKNLEDNLQSLATRLAPIYKQYA**
**PVAYQNQVEYENVARECRLGSKEGRPFSGVTACLDFCAHPHRDIHNMNNGSTVVCTLTREDNRSLGVI**
**PQDEQLHVLPLYKLSDTDEFGSKEGMEAKIKSGAIEVLAPRRKKRTCFTQPVPRSGKKRAAMMTEVLA**
**HKIRAVEKKPIPRIKRKNNSTTTNNSKPSSLPTLGSNTETVQPEVKSETEPHFILKSSDNTKTYSLMPSAP**
**HPVKEASPGFSWSPKTASATPAPLKNDATASCGFSERSSTPHCTMPSGRLSGANAAAADGPGISQLG**
**EVAPLPTLSAPVMEPLINSEPSTGVTEPLTPHQPNHQPSFLTSPQDLASSPMEEDEQHSEADEPPSDEP**
**LSDDPLSPAEEKLPHIDEYWSDSEHIFLDANIGGVAIAPAHGSVLIECARRELHATTPVEHPNRNHPTRL**
**SLVFYQHKNLNKPQHGFELNKIKFEAKEAKNKKMKASEQKDQAANEGPEQSSEVNELNQIPSHKALTL**
**THDNVVTVSPYALTHVAGPYNHWV**ID

Name: **3xFlag-PUFa-TET1(CD)**

**Keys:** NLS, PUFa, **TET1(1418-2136), 3xFlag tag**

M**DYKDHDGDYKDHDIDYKDDDDK**IDGGGGSDPKKKRKVDPKKKRKVDPKKKRKVGSTGSRNDGGGGS
GGGGSGGGGSGRAGILPPKKKRKVSRGRSRLLEDFRNNRYPNLQLREIAGHIMEFSQDQHGSRFIQLKL
ERATPAERQLVFNEILQAAYQLMVDVFGNYVIQKFFEFGSLEQKLALAERIRGHVLSLALQMYGSRVIEKA
LEFIPSDQQNEMVRELDGHVLKCVKDQNGNHVVQKCIECVQPQSLQFIIDAFKGQVFALSTHPYGCRVIQ
RILEHCLPDQTLPILEELHQHTEQLVQDQYGNYVIQHVLEHGRPEDKSKIVAEIRGNVLVLSQHKFASNVV
EKCVTHASRTERAVLIDEVCTMNDGPHSALYTMMKDQYANYVVQKMIDVAEPGQRKIVMHKIRPHIATLR
KYTYGKHILAKLEKYYMKNGVDLGDPKKKRKVDPKKKRKVGGRGGGGSGGGGSGGGGSGPA**ELPTCS**
**CLDRVIQKDKGPYYTHLGAGPSVAAVREIMENRYGQKGNAIRIEIVVYTGKEGKSSHGCPIAKWVLRRS**
**SDEEKVLCLVRQRTGHHCPTAVMVVLIMVWDGIPLPMADRLYTELTENLKSYNGHPTDRRCTLNENRT**
**CTCQGIDPETCGASFSFGCSWSMYFNGCKFGRSPSPRRFRIDPSSPLHEKNLEDNLQSLATRLAPIYKQ**
**YAPVAYQNQVEYENVARECRLGSKEGRPFSGVTACLDFCAHPHRDIHNMNNGSTVVCTLTREDNRSL**
**GVIPQDEQLHVLPLYKLSDTDEFGSKEGMEAKIKSGAIEVLAPRRKKRTCFTQPVPRSGKKRAAMMTE**
**VLAHKIRAVEKKPIPRIKRKNNSTTTNNSKPSSLPTLGSNTETVQPEVKSETEPHFILKSSDNTKTYSLMP**
**SAPHPVKEASPGFSWSPKTASATPAPLKNDATASCGFSERSSTPHCTMPSGRLSGANAAAADGPGIS**
**QLGEVAPLPTLSAPVMEPLINSEPSTGVTEPLTPHQPNHQPSFLTSPQDLASSPMEEDEQHSEADEPPS**
**DEPLSDDPLSPAEEKLPHIDEYWSDSEHIFLDANIGGVAIAPAHGSVLIECARRELHATTPVEHPNRNHP**
**TRLSLVFYQHKNLNKPQHGFELNKIKFEAKEAKNKKMKASEQKDQAANEGPEQSSEVNELNQIPSHKA**
**LTLTHDNVVTVSPYALTHVAGPYNHWV**ID

Name: **3xFlag-hDAGG45-PUFa-TET1(CD)**

**Keys:** NLS, PUFa, GADD45A, **TET1(1418-2136), 3xFlag tag**

**DYKDHDGDYKDHDIDYKDDDDKID**GGGGSSGAATMTLEEFSAGEQKTERMDKVGDALEEVLSKALSQR
TITVGVYEAAKLLNVDPDNVVLCLLAADEDDDRDVALQIHFTLIQAFCCENDINILRVSNPGRLAELLLLETD
AGPAASEGAEQPPDLHCVLVTNPHSSQWKDPALSQLICFCRESRYMDQWVPVINLPERSRTGAATMIDG
GGGSDPKKKRKVDPKKKRKVDPKKKRKVGSTGSRNDGGGGSGGGGSGGGGSGRAGILPPKKKRKVS
RGRSRLLEDFRNNRYPNLQLREIAGHIMEFSQDQHGSRFIQLKLERATPAERQLVFNEILQAAYQLMVDV
FGNYVIQKFFEFGSLEQKLALAERIRGHVLSLALQMYGSRVIEKALEFIPSDQQNEMVRELDGHVLKCVKD
QNGNHVVQKCIECVQPQSLQFIIDAFKGQVFALSTHPYGCRVIQRILEHCLPDQTLPILEELHQHTEQLVQ
DQYGNYVIQHVLEHGRPEDKSKIVAEIRGNVLVLSQHKFASNVVEKCVTHASRTERAVLIDEVCTMNDGP
HSALYTMMKDQYANYVVQKMIDVAEPGQRKIVMHKIRPHIATLRKYTYGKHILAKLEKYYMKNGVDLGDP
KKKRKVDPKKKRKVGGRGGGGSGGGGSGGGGSGPA**ELPTCSCLDRVIQKDKGPYYTHLGAGPSVAA**
**VREIMENRYGQKGNAIRIEIVVYTGKEGKSSHGCPIAKWVLRRSSDEEKVLCLVRQRTGHHCPTAVMVV**

**LIMVWDGIPLPMADRLYTELTENLKSYNGHPTDRRCTLNENRTCTCQGIDPETCGASFSFGCSWSMYF NGCKFGRSPSPRRFRIDPSSPLHEKNLEDNLQSLATRLAPIYKQYAPVAYQNQVEYENVARECRLGSK EGRPFSGVTACLDFCAHPHRDIHNMNNGSTVVCTLTREDNRSLGVIPQDEQLHVLPLYKLSDTDEFGSK EGMEAKIKSGAIEVLAPRRKKRTCFTQPVPRSGKKRAAMMTEVLAHKIRAVEKKPIPRIKRKNNSTTTN NSKPSSLPTLGSNTETVQPEVKSETEPHFILKSSDNTKTYSLMPSAPHPVKEASPGFSWSPKTASATPA PLKNDATASCGFSERSSTPHCTMPSGRLSGANAAAADGPGISQLGEVAPLPTLSAPVMEPLINSEPST GVTEPLTPHQPNHQPSFLTSPQDLASSPMEEDEQHSEADEPPSDEPLSDDPLSPAEEKLPHIDEYWSD SEHIFLDANIGGVAIAPAHGSVLIECARRELHATTPVEHPNRNHPTRLSLVFYQHKNLNKPQHGFELNKI KFEAKEAKNKKMKASEQKDQAANEGPEQSSEVNELNQIPSHKALTLTHDNVVTVSPYALTHVAGPYN HWV**ID
